# Genotypic and phenotypic consequences of domestication in dogs

**DOI:** 10.1101/2024.05.01.592072

**Authors:** Sweetalana, Shirin Nataneli, Shengmiao Huang, Jazlyn A Mooney, Zachary A Szpiech

## Abstract

Runs of homozygosity (ROH) are genomic regions that arise when two copies of identical haplotypes are inherited from a shared common ancestor. In this study, we leverage ROH to identify associations between genetic diversity and non-disease phenotypes in *Canis lupus familiaris* (dogs). We find significant association between the ROH inbreeding coefficient (F_ROH_) and several phenotypic traits. These traits include height, weight, lifespan, muscled, white coloring of the head and chest, furnishings, and fur length. After correcting for population structure, we identified more than 45 genes across the examined quantitative traits that exceed the threshold for suggestive significance. We observe distinct distributions of inbreeding and elevated levels of long ROH in modern breed dogs compared to more ancient breeds, which aligns with breeding practices during Victorian era breed establishment. Our results highlight the impact of non-additive variation and of polygenicity on complex quantitative phenotypes in dogs due to domestication and the breed formation bottleneck.

## Background

For over a century, scholars and dog-enthusiasts alike have sought to unravel the complex evolutionary history of man’s best friend [1]. *Canis lupus familiaris* (dogs) have intrigued researchers, across scientific fields, due to their close knit ties to our species [2] and unique genetic features [3–5]. While the exact origins of domestication remain elusive, understanding the impact of this process on the phenotypic variance and evolutionary history of the species is important.

In the 19th century, Darwin suggested that the broad range of features exhibited in modern breeds is conducive to a descendancy from multiple canid species, including wolves and jackals. In his accounts on variation, he argues that a single ancestral origin from the gray wolf (*Canis lupus*) seems to contradict the historical and archeological evidence available. Given that many wild species of the genus Canis have the potential to interbreed, Darwin’s proposal held fast for many years [1]. Advances in genome technology provided scientists with a new approach to studying dog ancestry, offering insights that complement and, in some cases, challenge the conclusions drawn from archaeological and historical evidence [6]. One commonly used approach is mitochondrial DNA (mtDNA) analysis which allows researchers to trace maternal ancestry. Using samples collected from dogs, wolves, and several other wild canid species, researchers found the similarity between dogs and the gray wolf to be significantly higher than that of dogs and any other wild canid species [7, 8]. In recent years, whole genome sequence (WGS) data has increasingly focused on the wolf-dog relationship, further supporting this finding and making it a primary area of investigation in contemporary studies [4, 9, 10].

While most agree that dogs were the first domesticated species, establishing their spatial and temporal roots has been a challenge [11]. The earliest depictions of human-canine ties come by means of cave paintings discovered in Saudi Arabia [12]. This cooperative hunting scene is believed to be 8000 to 9000 years old, but evidence gathered from burial sites predict an earlier origin of domestication [13]. The first dates back approximately 12,000 years ago to the Natufian in Northern Israel, where they discovered three sets of dog remains buried with a human [14, 15]. Additionally, a dog-like mandible was recovered at a burial site in Bonn-Oberkassel, Germany. This Northern European sample is believed to be about 14,000 years old [16, 17], however genetic work has added an additional layer of evidence for an even older wolf-dog divergence.

Genomic work, based on mtDNA and WGS data, currently offers two potential explanations for the origins of domestication. The first is that early dogs were a result of a single domestication event from gray wolves in East Asia between 16,000 and 100,000 years ago. The hypothesis stems from the limited haplotypic diversity in dogs, which may suggest a genetic bottleneck where domesticated dogs originated from a small-closely-related population of gray wolves [7]. Additionally, high levels of genetic diversity from the region south of the Yangtze River, indicate an older divergence than other areas in Europe and Asia. [18–20].

The second genomic narrative centers upon the theory that haplogroups entered the dog lineage at different times and places. Using nuclear genomic single nucleotide polymorphism (SNP) analysis, one study found evidence for allele sharing in Asian dog breeds and Asian wolves as well as European dog breeds and European wolves [21]. This work suggests that there were several founder events derived from separate wolf populations. Further WGS work has documented significant genetic differences in Eastern and Western Eurasian dog populations, providing further support for multiple domestication events [22, 23]. Despite these findings, the origins of domestication in dogs remain a hot topic of debate, warranting the need for additional studies exploring the complexities of their history.

While the early evolution of dogs from wolves marked a significant shift between species, the more recent history of established dog breeds has shaped evolution within the species itself. During the Victorian era, the selective breeding of aesthetically desirable traits led to the emergence of hundreds of new breeds. Breed establishment typically involved a limited number of founding members, resulting in high levels of inbreeding and dramatic loss of genetic diversity [3, 24–28]. Founder effects, such as this, restrict the gene pool and result in long stretches of DNA sharing between individuals. Shared segments derived from a single common ancestor are said to be identical-by-descent (IBD). Segments of the genome inherited IBD, when found within an individual, are commonly referred to as runs of homozygosity (ROH). Artificial selection for extreme phenotypes during breed formation made much of the genome homozygous and simplified complex trait architecture in dogs, which has facilitated the study of phenotypic associations between breeds [3, 29, 30].

Studies of ROH in humans have revealed associations with numerous complex disease and non-disease phenotypes, which provide valuable insight on the impact of genetic diversity on health and evolutionary processes [31–36]. Similar work has been done in dogs, providing evidence of association between ROH and several disease phenotypes [3, 30, 37]. Additionally, previous studies have explored the influence of genetic variants on quantitative traits such as; leg length, wrinkled skin, coat color, hair length, and skeletal shape [38–42]. Despite these advances, associations with complex non-disease phenotypes in dogs are relatively unexplored [43].

In this study, we investigate the relationship between the genomic consequences of domestication and complex (non-disease) trait architecture by using the distribution of ROH to provide insights on complex phenotypic architecture. To examine the relationship between ROH and non-disease phenotypes in dogs, we analyzed 556 canid whole-genome sequences and characterized the genomic distribution of ROH. After accounting for breed structure, we test for associations between the total ROH coverage of the genome (as measured by F_ROH_) among 13 breed groups with 13 phenotypes. Finally, we perform an ROH-mapping genome-wide association study and identify multiple genetic variants associated with phenotype, namely height, weight, and lifespan.

## Results

### ROH distribution

ROH were categorized into 6 length classes: Class A (0.5 - 1 Mbp), Class B (1 - 2 Mbp), Class C (2 - 3 Mbp), Class D (3 - 4 Mbp), Class E (4 - 5 Mbp), and Class F (>5 Mbp). For our 556 samples, the mean number of ROH (n_ROH_) was 37.06 and the mean length of ROH (s_ROH_) was 94.17 Mbp. These parameters were also computed for each ROH length class: Class A (n_ROH_ = 99.44; s_ROH_ = 68.39 Mbp), Class B (n_ROH_ = 50.29; s_ROH_ = 70.36 Mbp), Class C (n_ROH_ = 21.60; s_ROH_ = 52.81 Mbp), Class D (n_ROH_ = 13.07; s_ROH_ = 45.23 Mbp), Class E (n_ROH_ = 9.21; s_ROH_ =41.15 Mbp), and Class F (n_ROH_ = 28.77; s_ROH_ = 287.05 Mbp) (Table 1, Table 2).

**Table 1:**
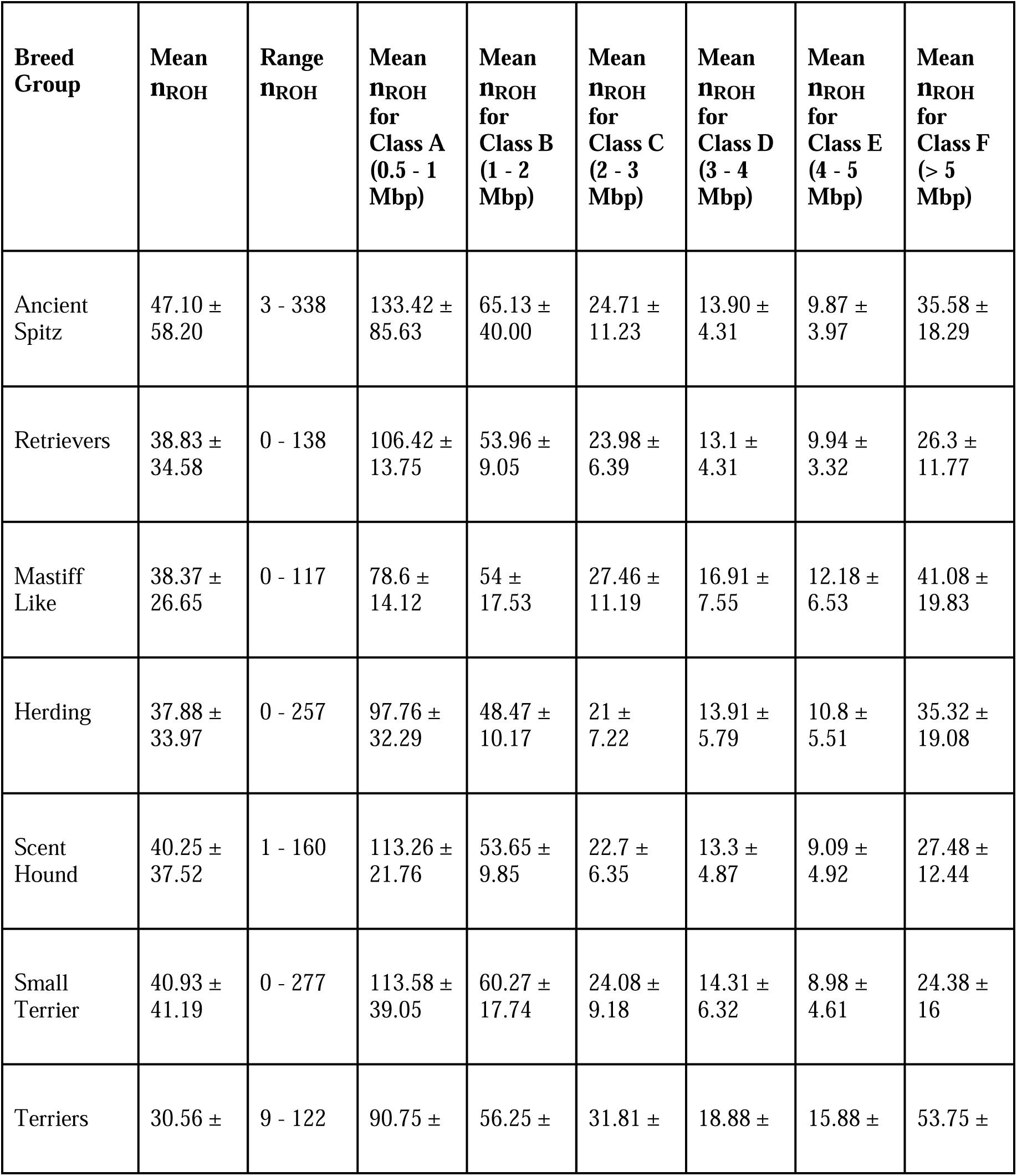
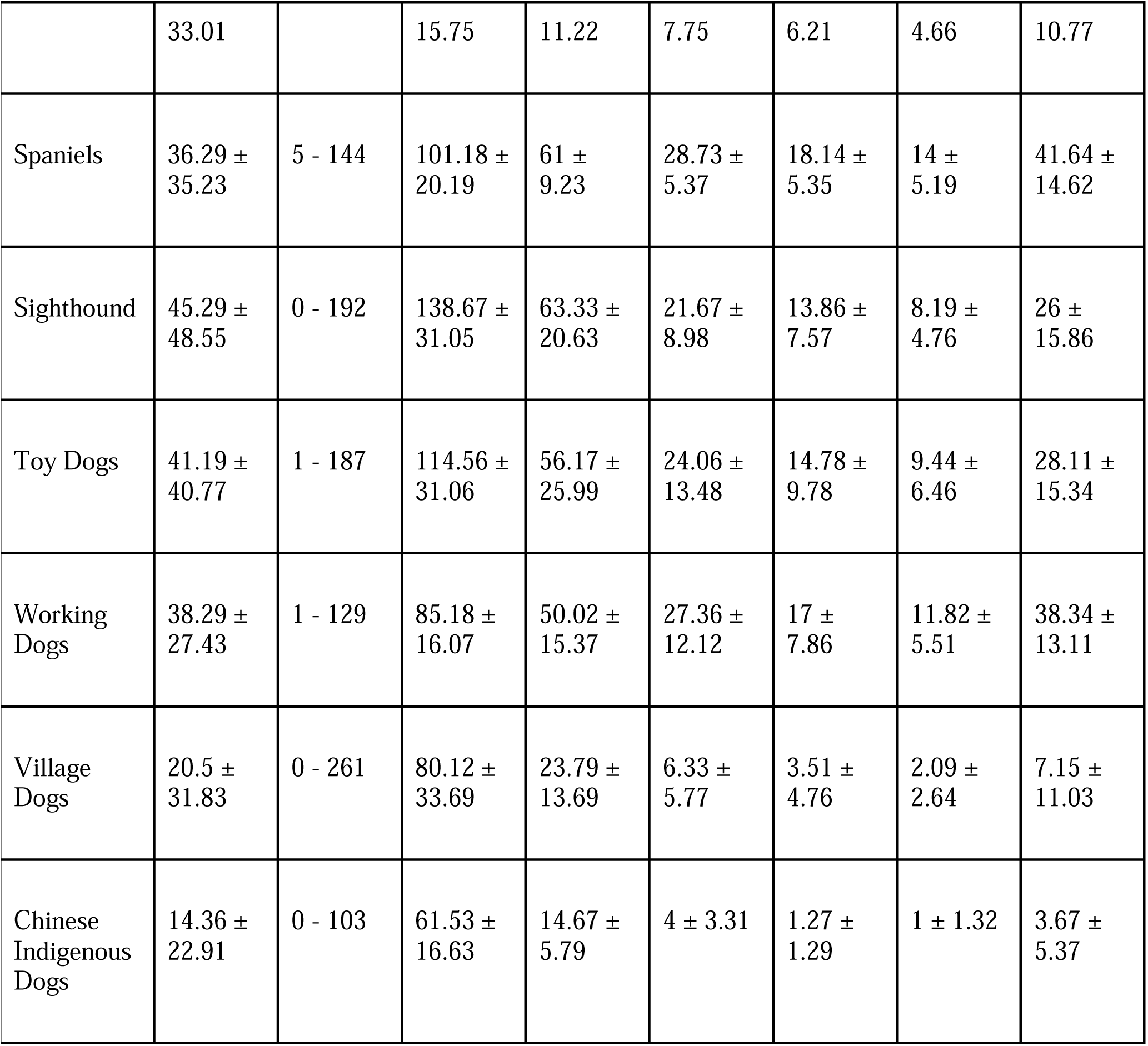
n_ROH_ across all breed groups and ROH length classification. This table details the range and mean ± standard deviation of the number of runs of homozygosity for each breed group and ROH length classification.

**Table 2:** s_ROH_ across all breed groups and ROH length classification. The table contains the range and mean ± standard deviation of the sum of lengths of runs of homozygosity for each breed group and ROH length classification.

| **Breed Group** | **Mean SROH (Mbp)** | **Range SROH (Mbp)** | **Mean SROH for Class A (0.5 - 1 Mbp)** | **Mean SROH for Class B (1 - 2 Mbp)** | **Mean SROH for Class C (2 - 3 Mbp)** | **Mean SROH for Class D (3 - 4 Mbp)** | **Mean SROH for Class E (4 - 5 Mbp)** | **Mean SROH for Class F (>5 Mbp)** |
| --- | --- | --- | --- | --- | --- | --- | --- | --- |
| Ancient Spitz | 119.56 ±<br>162.09 | 13.73 -<br>1113.00 | 92.29 ±<br>59.41 | 90.67 ±<br>54.30 | 60.33 ±<br>27.20 | 48.30 ± | 44.14 ±<br>17.51 | 381.60 ±<br>255.7 |
| Retrievers | 89.28 ±<br>86.63 | 0 -<br>598.02 | 72.96 ±<br>9.44 | 75.50 ±<br>13.25 | 57.37 ±<br>16.00 | 45.22 ±<br>15.07 | 44.47 ±<br>14.90 | 240.17 ±<br>125.98 |
| Mastiff Like | 126.47 ±<br>176.91 | 0 -<br>1029.28 | 54.77 ±<br>9.71 | 76.97 ±<br>25.95 | 67.21 ±<br>27.55 | 58.57 ±<br>25.92 | 54.38 ±<br>29.38 | 446.89 ±<br>247.25 |
| Herding | 105.25 ±<br>139.34 | 0 -<br>861.74 | 67.24 ±<br>21.71 | 67.53 ±<br>14.78 | 51.27 ±<br>17.64 | 48.23 ±<br>20.12 | 48.26 ±<br>24.73 | 348.94 ±<br>206.94 |
| Scent Hound | 99.19 ±<br>113.14 | 4.59 -<br>946.54 | 77.78 ±<br>14.73 | 74.84 ±<br>15.22 | 55.26 ±<br>15.67 | 45.90 ±<br>16.82 | 40.62 ±<br>21.78 | 300.75 ±<br>159.58 |
| Small Terrier | 89.41 ±<br>96.13 | 0 -<br>690.33 | 78.20 ±<br>26.38 | 84.24 ±<br>24.77 | 59.07 ±<br>22.76 | 49.63 ±<br>22.10 | 40.18 ±<br>20.51 | 225.15 ±<br>170.89 |
| Terriers | 75.86 ±<br>116.57 | 28.77 -<br>951.29 | 63.93 ±<br>10.90 | 79.69 ±<br>16.00 | 77.98 ±<br>19.72 | 65.84 ±<br>21.84 | 71.06 ±<br>20.79 | 541.71 ±<br>161.42 |
| Spaniels | 94.03 ±<br>127.54 | 23.23 -<br>856.91 | 70.13 ±<br>12.83 | 85.21 ±<br>12.79 | 70.39 ±<br>13.56 | 62.52 ±<br>18.65 | 62.55 ±<br>23.16 | 427.24 ±<br>190.19 |
| Sighthound | 94.65 ±<br>96.81 | 0 -<br>591.60 | 95.24 ±<br>21.50 | 86.87 ±<br>29.21 | 53.04 ±<br>22.07 | 47.98 ±<br>26.13 | 36.54 ±<br>21.32 | 248.26 ±<br>149.49 |
| Toy Dogs | 97.01 ±<br>107.57 | 4.12 -<br>602.43 | 78.92 ±<br>22.17 | 79.01 ±<br>36.61 | 58.31 ±<br>33.13 | 50.45 ±<br>33.72 | 42.33 ±<br>28.82 | 273.02 ±<br>162.01 |
| Working<br>Dogs | 112.89 ±<br>130.13 | 4.74 -<br>658.42 | 58.67 ±<br>11.14 | 70.67 ±<br>22.08 | 66.54 ±<br>29.19 | 58.83 ±<br>27.51 | 52.76 ±<br>24.70 | 369.85 ±<br>139.08 |
| Village<br>Dogs | 33.24 ±<br>59.75 | 0 -<br>554.84 | 54.07 ±<br>23.09 | 32.65 ±<br>19.59 | 15.30 ±<br>13.97 | 12.04 ±<br>16.34 | 9.35 ±<br>11.72 | 76.00 ±<br>127.67 |
| Chinese<br>Indigenous<br>Dogs | 19.82 ±<br>35.06 | 0 -<br>309.27 | 41.03 ±<br>11.34 | 20.00 ±<br>7.64 | 9.53 ±<br>7.68 | 4.27 ±<br>4.24 | 4.41 ±<br>5.87 | 39.67 ±<br>75.16 |

Domestic dog clades showed a distinct pattern of genetic diversity, with higher levels of long ROH (>5 Mbp) when compared with Chinese indigenous dogs (light green) and village dogs (royal blue) (Figs. 1 & 2). The highest total n_ROH_ (641) was observed in Dingo01 and the highest total s_ROH_ (1643.98 Mbp) was observed in NGSD2, both individuals were from the Ancient Spitz breed group (Table S1). Terriers had the smallest mean n_ROH_ and the smallest mean s_ROH_ when compared with the remaining 11 breed groups (Tables 1 & 2).

**Fig. 1:**
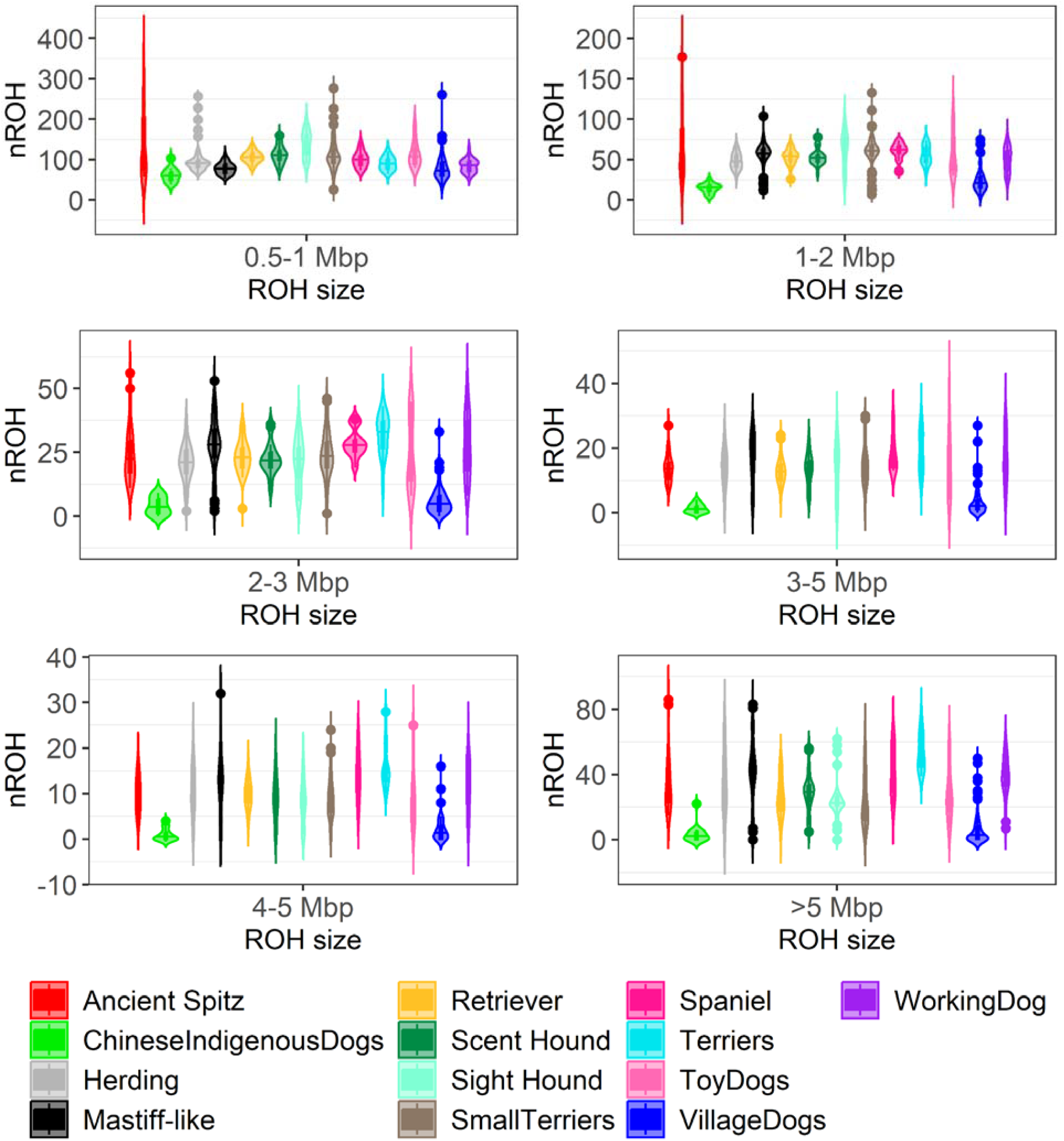
Relationship between the number of runs of homozygosity (n_ROH_) and ROH size class in Mbp for all breed groups. The x-axis represents the ROH size in Mbp. The y-axis represents the number of ROH. Each violin plot depicts the distribution of the total number of ROH within length classes 0.5-1 Mbp, 1-2 Mbp, 2-3 Mbp, 3-4 Mbp, 4-5 Mbp and >5 Mbp for each breed group.

**Fig. 2:**
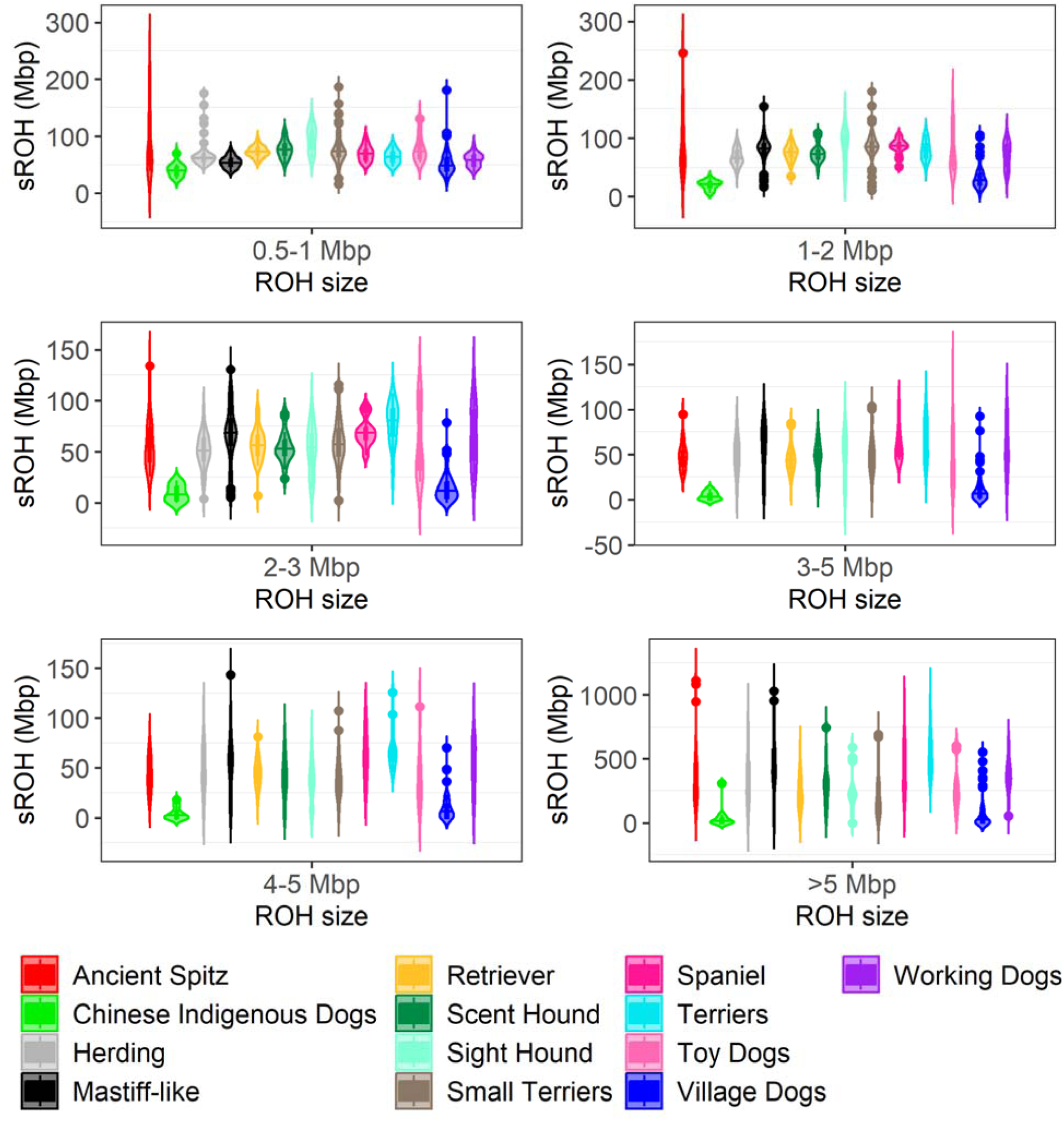
Relationship between the sum of lengths of runs of homozygosity (s_ROH_) and ROH size classes. The x-axis represents our defined ROH size classes in Mbp. The y-axis represents s_ROH_, which also has units of Mbp. Each violin plot depicts the distribution of the sum of ROH within length ranges 0.5-1 Mbp, 1-2 Mbp, 2-3 Mbp, 3-4 Mbp, 4-5 Mbp and >5 Mbp for each breed group.

Next, we explored the relationship between the total number of ROH (n_ROH_) and the total lengths of ROH (s_ROH_) (Fig. 3). We observed significantly higher mean values for both n_ROH_ and s_ROH_ for the domesticated breed dogs relative to the domesticated non-breed Chinese indigenous dogs and village dogs. Breed dogs also tended to fall off of the y=x line when n_ROH_ was compared to s_ROH_. The accumulation of long ROH within breed dogs in the recent time was most drastic in Terrier clade, where we also observed the largest F_ROH_ (Table 3). We observed the largest variance in nROH in the Small Terriers, which include breeds such as the Australian Terrier, Cairn Terrier, Jack Russell Terrier, Norwich Terrier, Scottish Terrier, Tibetan Terrier, West Highland, White Terrier, and Yorkshire Terrier. We also observed quite a bit of variance in Ancient Spitz, Village dogs, Herding, Mastiff-like, and Toy dogs. Conversely, Chinese Indigenous dogs, Terriers, Spaniels, Retrievers, and Scent Hound cluster together quite tightly. The Spaniels fall somewhere between these groups.

**Fig. 3:**
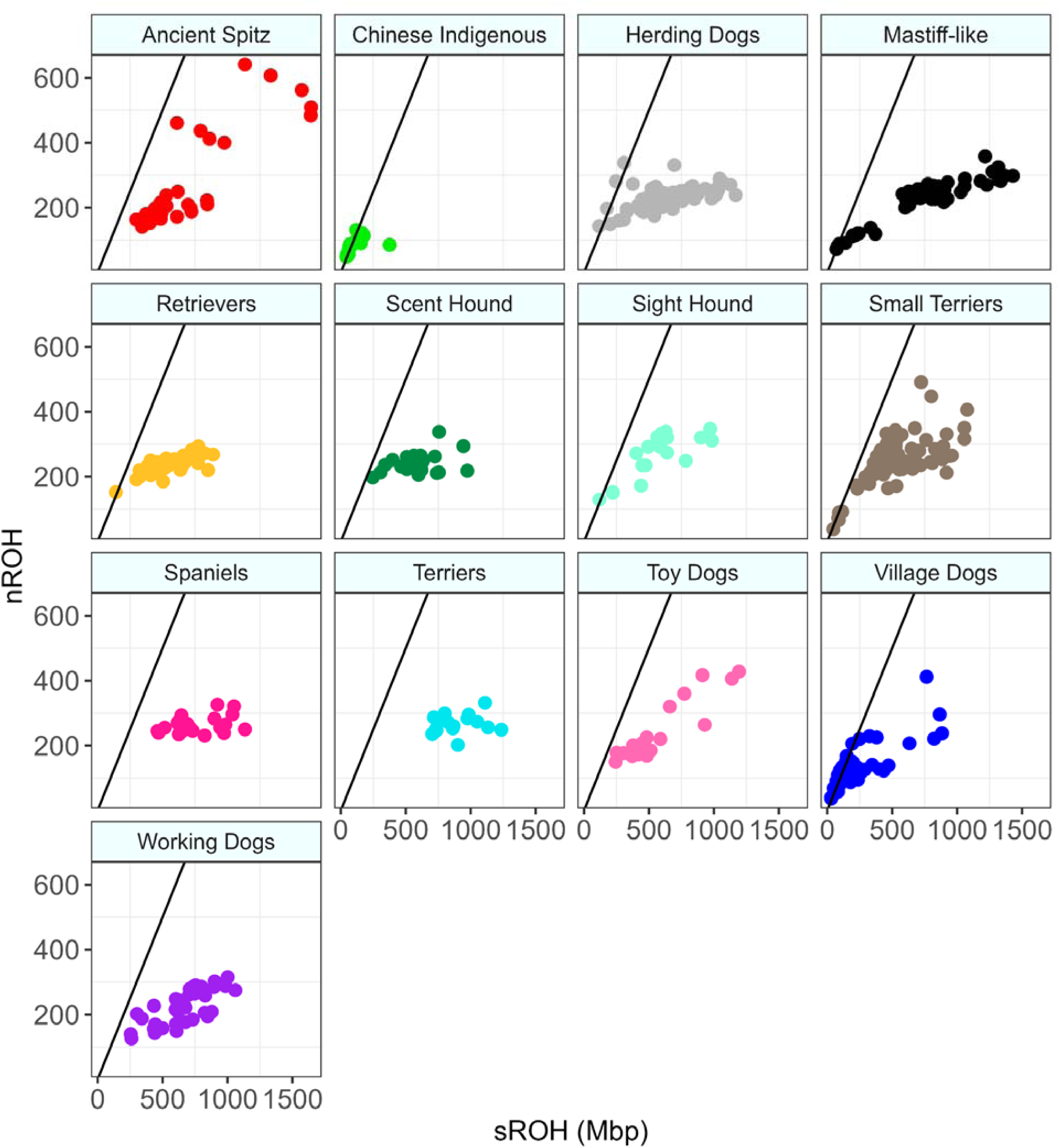
Relationship between the sum of lengths of runs of homozygosity (s_ROH_) and the number of runs of homozygosity (n_ROH_) for each breed group. The x-axis and the y-axis represent the sROH and nROH, respectively. The black line is y=x.

**Table 3:** F_ROH_ across all breed groups. The table represents the mean ± standard deviation of F_ROH_ for each breed group.

| Breed Group | Mean $F_{ROH}$ |
| --- | --- |
| Terriers | $0.41 \pm 0.07$ |
| Spaniels | $0.35 \pm 0.09$ |
| Mastiff-Like | $0.34 \pm 0.15$ |
| Ancient Spitz | $0.33 \pm 0.17$ |
| Working Dog | $0.31 \pm 0.09$ |
| Herding | $0.29 \pm 0.11$ |
| Scent Hound | $0.27 \pm 0.08$ |
| Toy Dogs | $0.26 \pm 0.13$ |
| Sight Hound | $0.26 \pm 0.10$ |
| Small Terrier | $0.24 \pm 0.10$ |
| Retriever | $0.24 \pm 0.07$ |
| Village Dogs | $0.09 \pm 0.08$ |
| Chinese Indigenous Dogs | $0.05 \pm 0.04$ |

We were also interested in whether any of ROH hotspots were shared across clades or between breed dogs and domesticated non-breed dog populations. We examined ROH coverage per site for each chromosome and plot sites where more than 50% of individuals within each clade have a ROH (Fig. S1-S16). There were clear hotspots within clades on every chromosome. Some of these hotspots corresponded with recombination rate, such as on chromosome 22, where we observed lower recombination rate in regions where there were shared ROH hotspots among all individuals. Conversely, on chromosome 12 we observed a high recombination rate, and very few shared ROH hotspots among individuals. Moreover, we observed very few shared ROH hotspots among different clades. One stand out was chromosome 28, where we observed a shared ROH hotspot among Ancient Spitz, Mastiff Like, Retrievers and Terriers (Fig. S13). Likewise on chromosome 32, we observed a shared ROH hotspot among Mastiff-Like, Scent Hound, Terriers and Toy Dogs (Fig. S14). The majority of hotspots were shared within a clade. This was best shown on chromosome 17, where we found multiple clade-specific ROH hotspots (Fig. S9). The same pattern was seen on chromosomes 31-38, with a few instances of sharing between Mastiff-like breeds and Terriers (Fig. S13 - S16). Lastly, we rarely observed any peaks for village dogs and Chinese Indigenous dogs (Fig. S3 - S16).

Next, we examined the level of inbreeding within clades by computing the inbreeding coefficient, F_ROH_, for each individual and each breed group (Tables 3 & S2). To account for the distribution of F_ROH_ across all breed groups, we computed mean F_ROH_ (Table 3). High levels of mean F_ROH_ were observed in breed dogs’ clades. F_ROH_ mean values in breed dogs ranged from 0.41 ± 0.07 in Terriers to 0.24 ± 0.07 in Retrievers. The lowest mean values were observed in non-breed dogs, namely Chinese Indigenous Dogs (0.05 ± 0.04) and Village Dogs (0.09 ± 0.08). This result was expected and provides support for breed dogs sharing a more recent common ancestor and more background relatedness relative to village or Chinese indigenous dogs [3].

### Association between F_ROH_ and phenotypic traits

Next, we wondered whether F_ROH_ could be used to detect non-additivity in quantitative (non-disease) phenotypes in dogs. Following an approach that was recently used in disease phenotypes in dogs [37] and quantitative and disease phenotypes in humans [32, 33, 44–48], we searched for non-additive effects by correlating F_ROH_ and 13 breed averaged phenotypes (bulky, drop ears, furnish, hairless, height, large ears, length of fur, lifespan, long legs, muscled, weight, white chest, and white head). The three quantitative phenotypes had significant associations with F_ROH_: breed average height (*β* = 0.31 and p = 4.05 × 10^-3^), breed average weight (*β* = 0.19 and p = 1.19 × 10^-2^), and breed average lifespan (*β* = -0.14 and p = 4.44 × 10^-2^) (Fig. 4). In breed dogs, we found that as F_ROH_ increased by 1%; breed average height increased by 0.192 cm, breed average weight increased by 0.14 kg, and breed average lifespan decreased by 0.014 years. The statistical analysis, including normalized coefficient effect size (either β or log odds), p-values, and confidence intervals for quantitative traits (breed average height, breed average weight, and breed average lifespan) are shown in Table S3. Significant associations with F_ROH_ were present in 5 out of the 10 remaining qualitative phenotypes: muscled (*β* = 7.96 and p = 9.68 × 10^-11^), white chest (*β* = 3.49 and p = 5.36 × 10^-5^), white head (*β* = 4.26 and p = 1.55 × 10^-4^), length of fur (*β* = -1.88 and p = 1.27 × 10^-2^), and furnish (*β* = -2.50 and p= 8.24 × 10^-3^) (Fig. 5, Table S4). Across all breed groups, as F_ROH_ increased, white chest, white head and muscled phenotypes also increased, whereas length of fur and furnish phenotypes decreased.

**Fig. 4:**
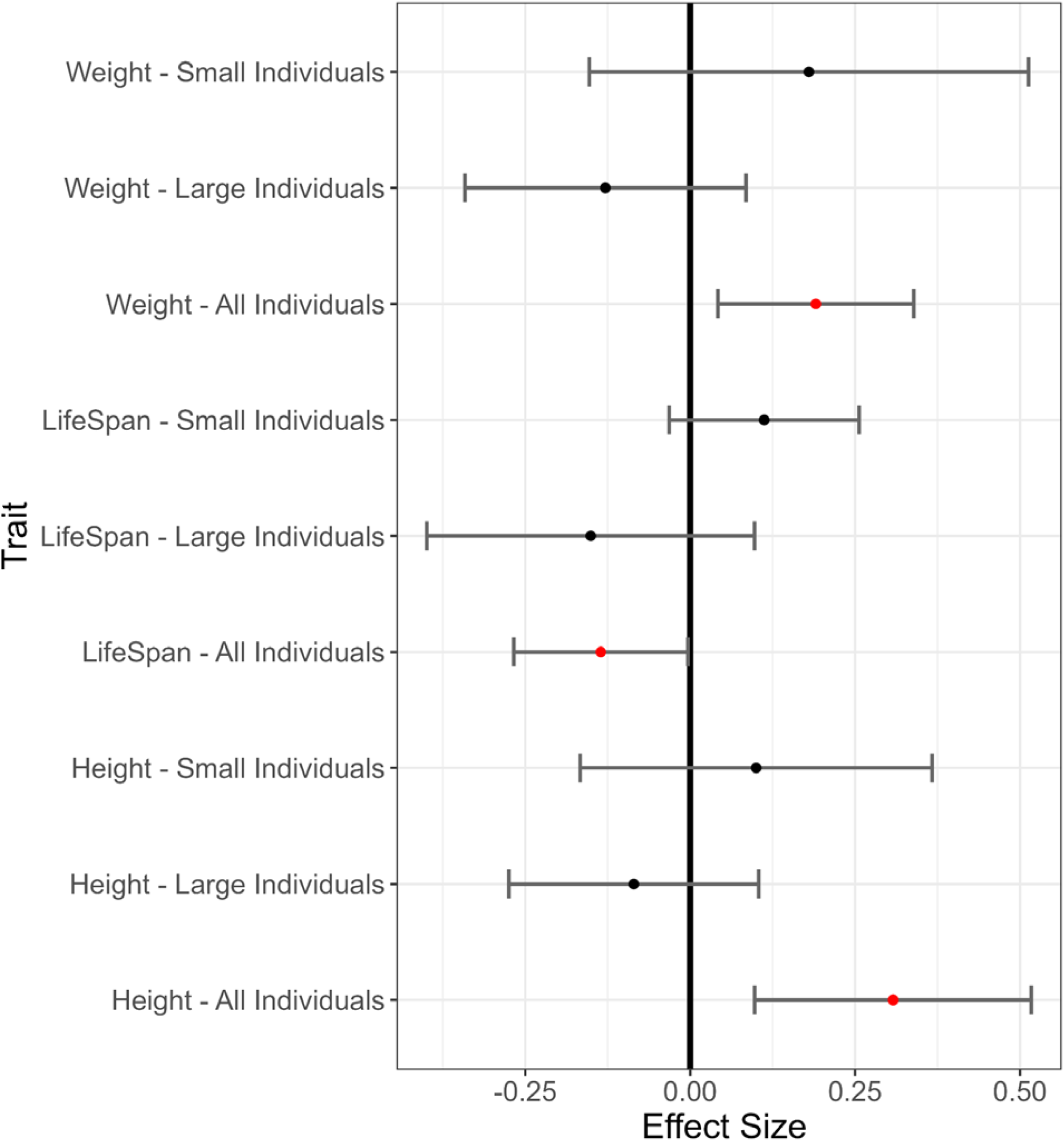
Relationship between effect size and breed averaged phenotypic traits. The y-axis represents the phenotypes for which the associations were tested. These are binned into 3 categories (small individuals, large individuals, and all individuals). The x-axis represents the normalized beta coefficients (β) for each trait. Significant effects of F_ROH_ on a phenotype (nominal p < 0.05) are indicated by a red point. An effect size larger than 0 indicates an increase of that trait value with respect to F_ROH,_ and less than 0 represents a decrease in that trait value.

**Fig. 5:**
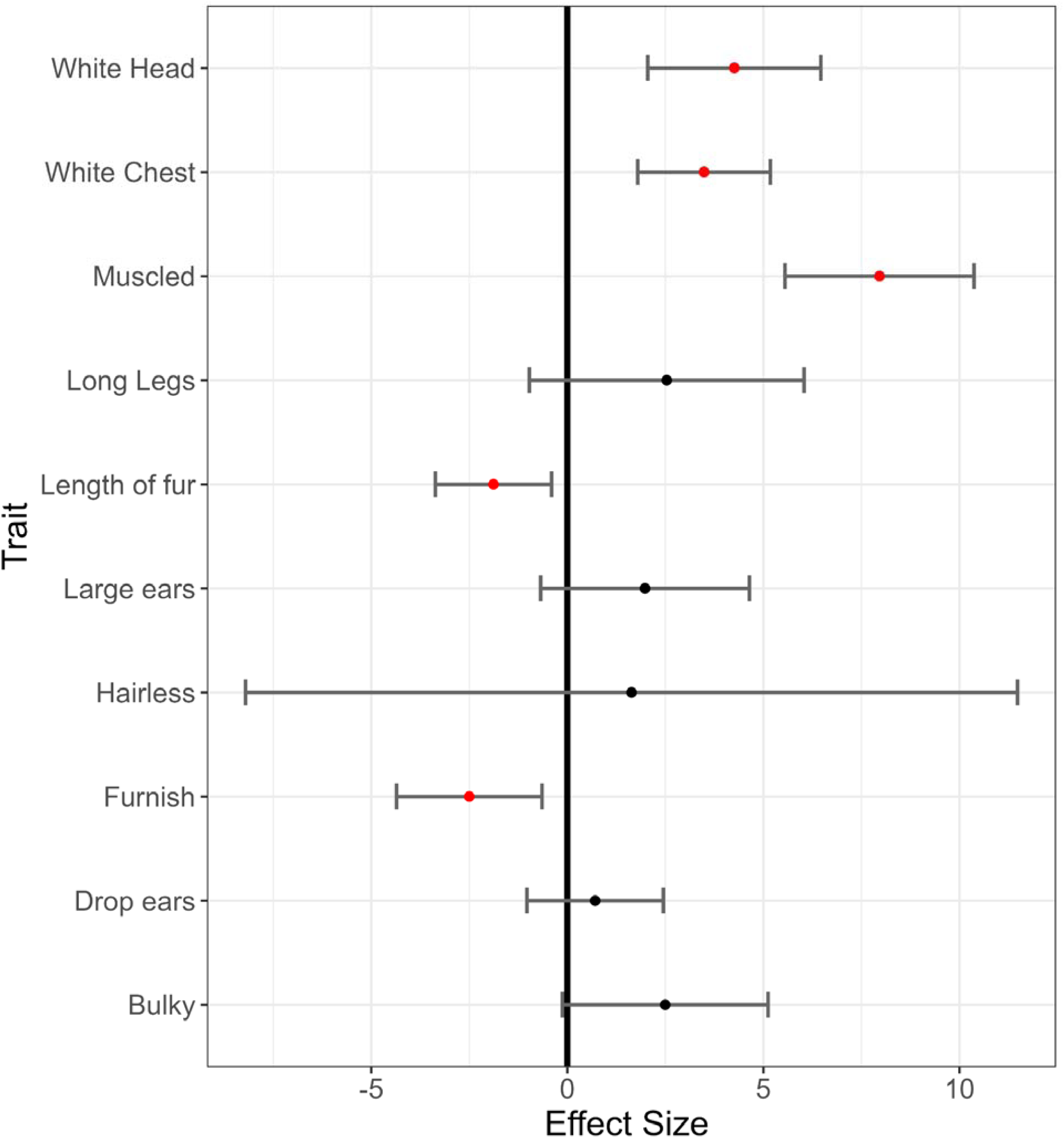
Relationship between effect size and categorical phenotypic traits. The y-axis represents the phenotypes for which the associations were tested. The x-axis represents the log odds for each trait. A significant effect of F_ROH_ on a phenotype (nominal p < 0.05) is indicated with a red point. An effect size larger than 0 indicates an increase of that trait with respect to F_ROH,_ and less than 0 represents a decrease in that trait.

### ROH-mapping GWAS

Finally, we sought to identify the ROH-associated genomic regions that influence our quantitative traits of interest (weight, lifespan, or height). Thus, we performed a GLMM-based GWAS (Fig. 6, Fig. 7, Fig. 8, and Fig. S17-S25) using presence/absence of ROH across individuals as the phenotype predictor for each locus. For exact significant threshold values, see Tables S4 - S5.

**Fig. 6.**
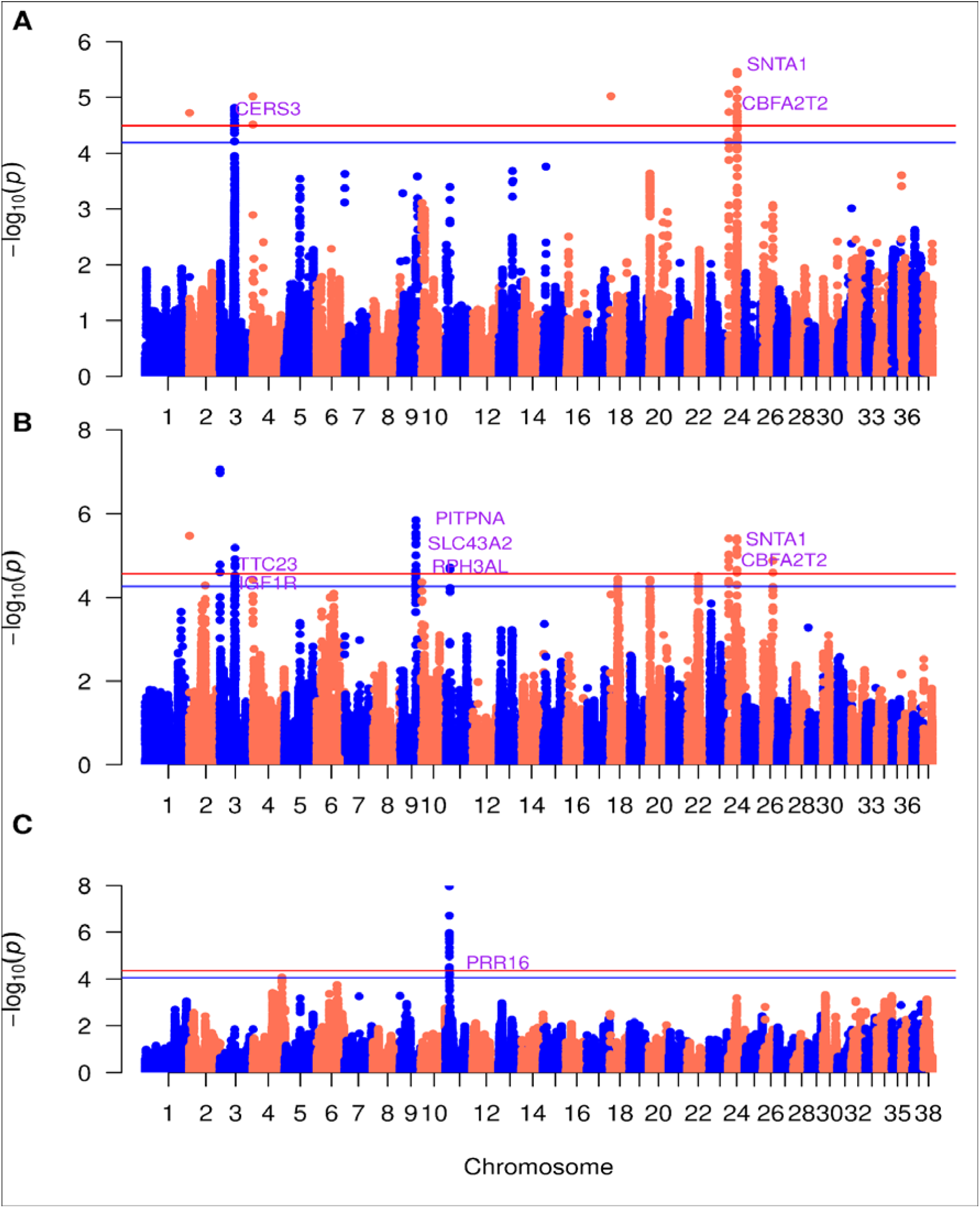
Manhattan plots for GLMM-based GWAS using presence/absence of ROH for body height. A. All individuals B. Small Individuals C. Large Individuals. The x-axis represents the genomic position. The y-axis represents the log10 base transformed p-values. Single nucleotide polymorphisms are represented by a single point. The red horizontal line indicates the genome wide significance (GWS) threshold, and the blue horizontal line indicates the suggestive wide significance (SWS) threshold. Only genes above the GWS threshold are labeled, with the exception of IGF1R. Marked genes are not drawn to scale.

**Fig. 7.**
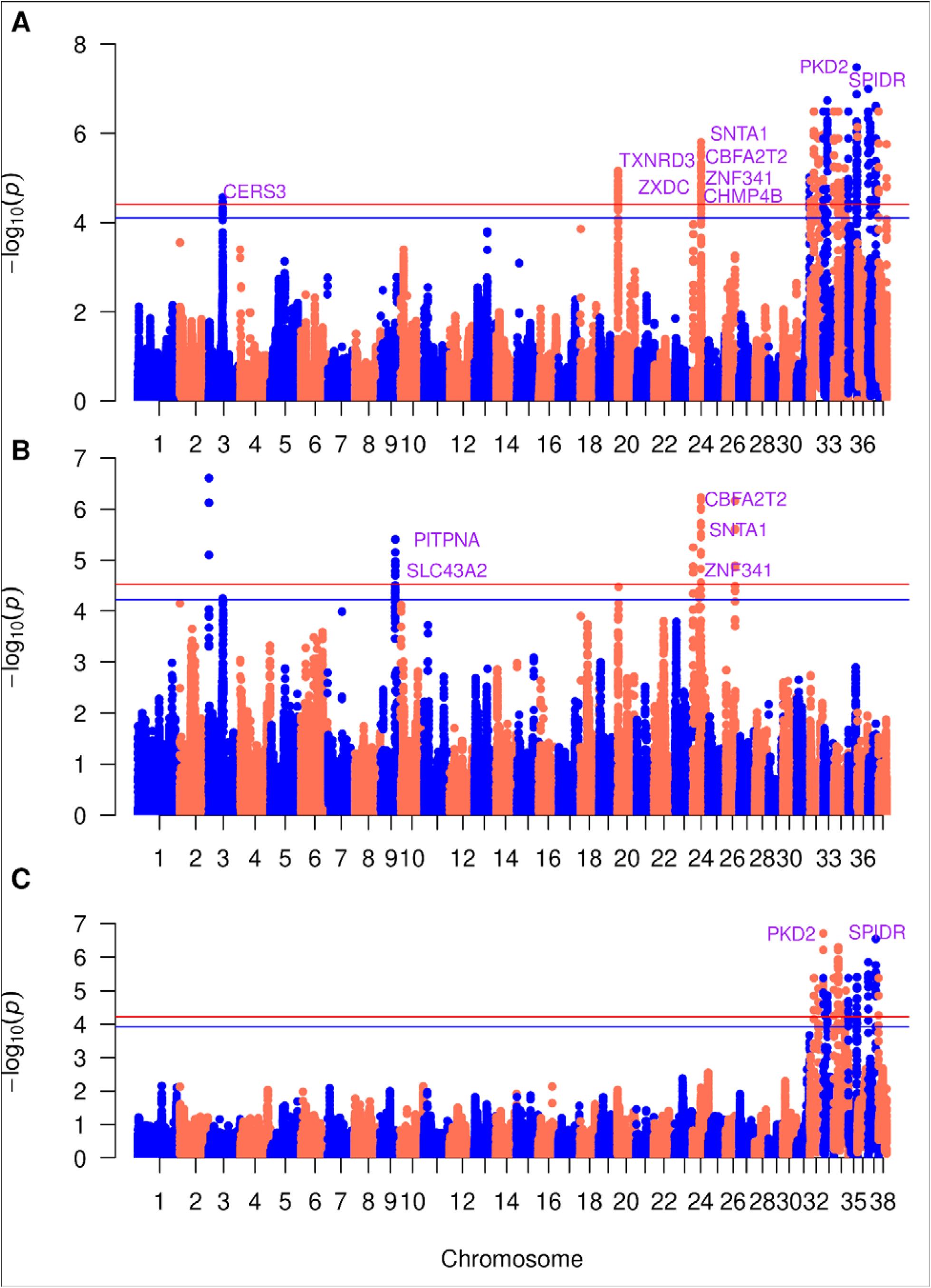
Manhattan plots for GLMM-based GWAS using presence/absence of ROH for body weight. A. All individuals B. Small Individuals C. Large Individuals. The x-axis represents the genomic position. The y-axis represents the log10 base transformed p-values. Single nucleotide polymorphisms are represented by a single point. The red horizontal line indicates the genome wide significance (GWS) threshold, and the blue horizontal line indicates the suggestive wide significance (SWS) threshold. Only genes above the GWS threshold are labeled. Marked genes are not drawn to scale.

**Fig. 8.**
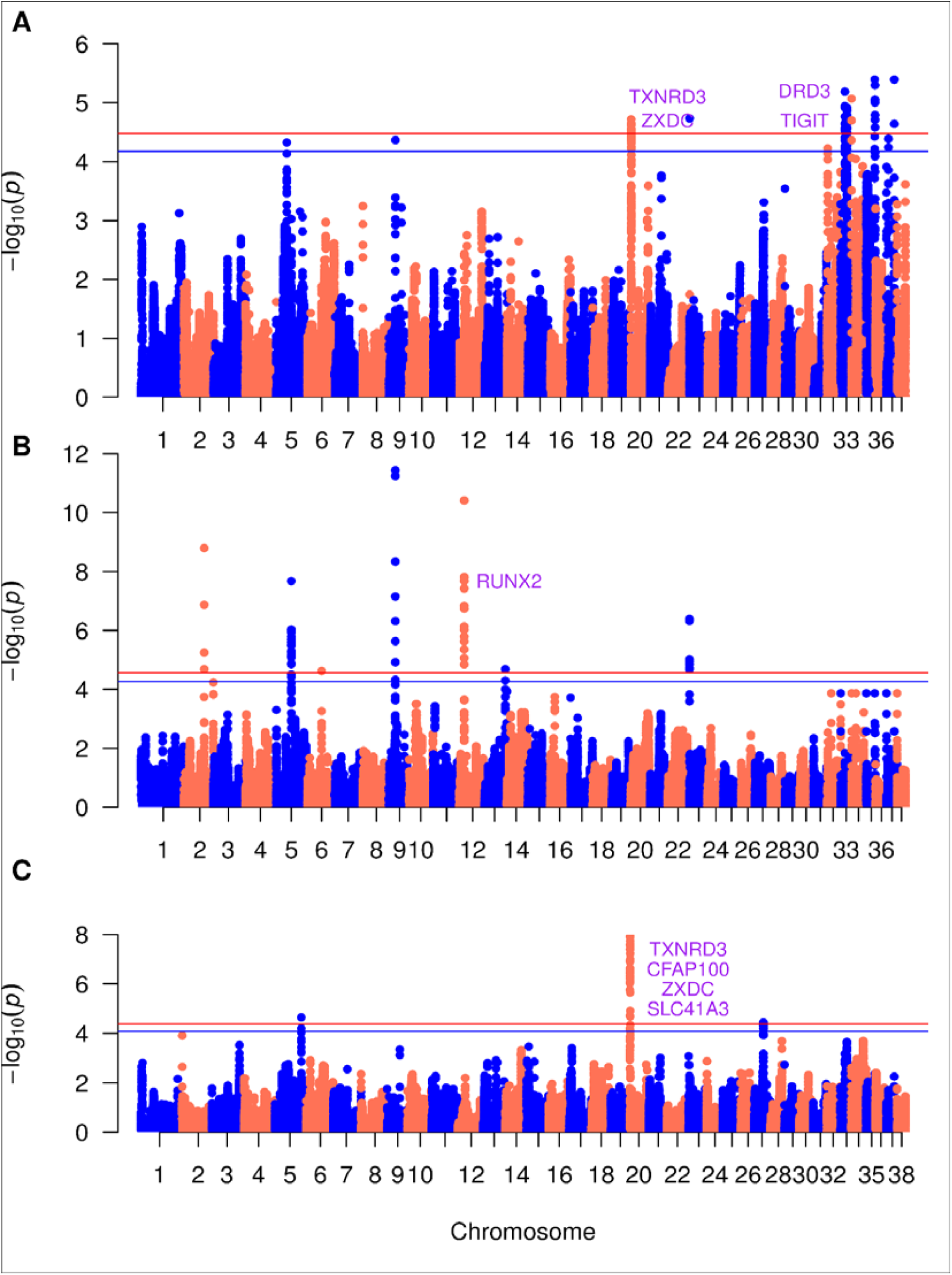
Manhattan plots for GLMM-based GWAS using presence/absence of ROH for lifespan. A. All individuals B. Small Individuals C. Large Individuals. The x-axis represents the genomic position. The y-axis represents the log10 base transformed p-values. Single nucleotide polymorphisms are represented by a single point. The red horizontal line indicates the genome wide significance (GWS) threshold, and the blue horizontal line indicates the suggestive wide significance (SWS) threshold. Only genes above the GWS threshold are labeled. Marked genes are not drawn to scale.

For breed average height, we found 27 SNPs above the Suggestive-Wide Significance (SWS) threshold and 18 SNPs above the Genome-Wide Significance (GWS) threshold when testing for all individuals in our sample (Fig. 6A). Notably, a single SNP on chromosome 24 (position 22647289) had the strongest associated genomic signal with height ( p = 3.46 × 10^-6^). This region corresponds with the gene *SNTA1*. Additionally, genes *CERS3* (7 SNPs) on chromosome 3, *CBFA2T2* (16 SNPs) on chromosome 24, and *SNTA1* (4 SNPs) on chromosome 24 were observed to be above the SWS threshold and contained multiple SNPs within their transcription regions (Table S6). SNPs linked to these three genes were also above the GWS threshold.

For small individuals, 34 SNPs were observed to be above the SWS threshold and 17 SNPs were observed to be above the GWS threshold when tested for association with height (Fig. 6B). A single SNP on chromosome 9 (position 45711401) had the strongest associated genomic signal (p = 1.44 × 10^-6^), which corresponds with the gene *PITPNA*. Across our small subgroup we identified additional genomic variants above the SWS threshold for weight: on chromosome 3, *CERS3* (3 SNPs), *IGF1R* (1 SNP), *LRRC28* (1 SNP), and *TTC23* (3 SNPs); on chromosome 9, *PITPNA* (4 SNPs), *SLC43A2* (2 SNPs), *VPS53* (2 SNPs), and *RPH3AL* (4 SNPs); on chromosome 18, *HGF* (2 SNPs); on chromosome 22, *COMMD6* (1 SNP); on chromosome 24, *CBFA2T2* (7 SNPs) and *SNTA1* (4 SNPs) (Table S6). Only SNPs associated with *TTC23*, *PITPNA*, *SLC43A2*, *RPH3AL*, CBFA2T2, and SNTA1 were above the GWS threshold. For large individuals, a single SNP on chromosome 11 (position 9943557), which is linked to *PRR16*, was observed to be above the SWS threshold for height (Fig. 6C). For an extended table containing all the observed genes associated with height refer to Table S6.

For breed average weight, 66 SNPs were observed above the SWS threshold and 45 SNPs above the GWS threshold, when including all individuals in our sample (Fig. 7A). Nearly half of the SNPs above the SWS threshold (30 SNPs) were found within genes on chromosome 24. A single SNP, on chromosome 32 (position 11413601) had the strongest association with weight (p =3.24 × 10^-7^). This was observed within *PKD2*, a gene that encodes for protein kinase D2. Across all samples we identified several other SNPS to be above the SWS threshold for weight: on chromosome 3, *CERS3* (7 SNPs); on chromosome 20, *ZXDC* (3 SNPs) and *TXNRD3* (1 SNP); on chromosome 24, *SNTA1* (5 SNPs), *CBFA2T2* (19 SNPs), *ZNF341* (2 SNPs), and *CHMP4B* (4 SNPs); on chromosome 32, *SEPT11* (1 SNP), *CCNI* (1 SNP), *PKD2* (11 SNPs), *SGMS2* (4 SNPs) and *CYP2U1* (3 SNPs); on chromosome 35, *SPIDR* (5 SNPs) (Table S7). The genes on chromosomes 3, 20, 24, and 35 all had SNPs above the GWS threshold. The only gene from chromosome 32 with SNPs above the GWS threshold was *PKD2*.

For small individuals, 31 SNPs were observed above the SWS threshold and 19 SNPS above the GWS threshold (Fig. 7B). Nearly a third of those above the SWS threshold (i.e. 20 SNPs) belong to chromosome 24. A single SNP on chromosome 24 (position 22660667) had the strongest association with weight ( p = 5.88 × 10^-7^) and is linked to the protein-coding gene *SNTA1* (Fig. 7B). For our subgroup of small individuals, we found SNPs above the SWS threshold for weight to be linked to the following genes: on chromosome 3, *CERS3* (1 SNP); on chromosome 9, *RPH3AL* (4 SNPs), PITPNA (4 SNPs), and SLC43A2 (2 SNPs); on chromosome 24, *ADAM33* (2 SNPs), *SIGLEC1* (2 SNPs), *HSPA12B* (1 SNP), *SNTA1* (4 SNPs), *CBFA2T2* (9 SNPs), *ZNF341* (1 SNP) and *CHMP4B* (1 SNP) (Table S7). SNPs on genes *PITPNA*, *SLC43A2*, *SNTA1*, *CBFA2T2*, and *ZNF341* were above the GWS threshold. In contrast, for the subgroup of large individuals, 10 SNPs were above the SWS threshold and 5 SNPs were above the GWS threshold (Fig. 7C). Only two genes reached significance in large individuals: on chromosome 32, *PKD2* (4 SNPs) and on chromosome 35, *SPIDR* (6 SNPs). All the SNPs linked to these two genes were above the SWS threshold for weight (Table S7). SNPs on both of these genes were also above the GWS threshold. For an extended table containing all the observed genes associated with weight refer to Table S7.

For breed average lifespan, 10 SNPs were observed above the SWS threshold and only 4 SNPs were above the GWS threshold when tested across all individuals (Fig. 8A). The strongest associated SNP (p= 2.21 × 10^-5^) was located on chromosome 20 (position 789204) within the gene *TXNRD3*. Across all samples we identified several other SNPS to be above the SWS threshold for lifespan: on chromosome 20, *ZXDC* (2 SNPs) and *TXNRD3* (1 SNP); on chromosome 32, *SEPT11* (1 SNP); on chromosome 33, *DRD3* (5 SNPs) and *TIGIT* (1 SNP) (Table S8). All, but *SEPT11*, had one SNP above the GWS threshold. For our subgroup of small individuals, we only found a gene on chromosome 12 *RUNX2* (4 SNPs), to be above the SWS threshold. (Fig. 8B and Table S8). For our subgroup of large individuals, we found 9 SNPs to be above the SWS threshold, all of which were also above the GWG threshold (Fig. 8C). The strongest associated SNP (p = 1.17 × 10^-8^) was located on chromosome 20 (position 789204) within the gene *TXNRD3.* All genes with SNPs above the significance thresholds (*TXNRD3* (1 SNP), *ZXDC* (3 SNPs), *CFAP100* (4 SNPs), and *SLC41A3* (1 SNP)) were located on chromosome 20 (Table S8). For an extended table containing all the observed genes associated with lifespan refer to Table S8.

## Conclusions

Our work provides in-depth analysis of patterns of runs of homozygosity (ROH) in domesticated breed dogs and non-breed domesticated village and Chinese Indigenous dogs, highlighting how the domestication and breed formation has shaped genetic diversity and trait architecture. We demonstrate that ROH, quantified as the total fraction of the genome within a run of homozygosity, F_ROH_, can be used to uncover traits that are not fully additive. We focus on ROH because these genomic segments reflect recent demography and inbreeding [49, 50]. For breed dogs, there is a large fraction of the genome within long ROH because of the small number of individuals used for breed establishment. These long ROH are a result of very recent parental relatedness and are inextricably linked to this species’ complex evolutionary, domestication, and breed formation history.

Previous research has shown that dogs originated from an isolated wolf population(s), and recent strong artificial selection drove breed emergence [3, 51]. Strong artificial selection during breed formation resulted in low genetic diversity and high homozygosity within breeds, alongside the large phenotypic variance between breeds [38]. Artificial selection, in the case of dogs, was achieved through inbreeding, and also led to the significant sharing of common genetic variants [52]. Artificial selection for specific phenotypic characteristics during breed establishment and subsequent breed standardization have resulted in homogeneity of both the phenotypic and genotypic variation within a breed. Further, when looking between breeds, previous work has shown that occasionally a dog with a desirable trait in one breed is also used to introduce the same phenotype in another breed, which creates a network of genetic relatedness through shared common haplotypes between breeds [53]. In sum, because artificial selection was often achieved through inbreeding, and breed standards reduced the effective population size of each breed, we expect both phenomena to be reflected through shared haplotypic information between breeds and distinct patterns of homozygosity within breeds.

For example, we show that domesticated breed dogs have unique patterns of ROH in their genome, and on average carry more of their genome within long ROH than domesticated non-breed village and Chinese indigenous dogs (Table 2). Conversely, the village dogs and Chinese indigenous dogs which did not experience selection from breed formation have the lowest values for mean ROH distribution. Taken together our results correspond with the higher degree of relatedness among all domestic breed group individuals. This was expected given the known high levels of inbreeding during breed establishment. Our results also highlight village and Chinese indigenous dogs as more outbred populations given lower ROH proportion when compared to breed dogs. Specifically, we observe lower levels of mean n_ROH_ and mean s_ROH_ in Chinese indigenous dogs. This is in line with the origins of the Chinese indigenous dogs, which is highlighted by relaxed trait selection during establishment of the population [54]. The relaxed selection may have resulted in overall higher levels of genetic diversity. Additionally, village dogs also harbor lower values for mean n_ROH_ and mean s_ROH_ (Table 1 and Table 2), consistent with them having not experienced an additional bottleneck during breed formation [37].

Turning to ROH hotspots, we found that there was some relationship with recombination rate per chromosome and the density of ROH hotspots, though recombination certainly does explain some of the observed hotspot and coldspots, it does not fully explain all of the patterns that we observed (Fig. S1 & S4). For example, on many chromosomes, we observe at least one ROH hotspot (across breed dogs and village dogs) in a region where recombination rates peaks. This is also consistent with the conclusions from previous studies which suggest recombination rates are not the only factor leading to ROH hotspots or coldspots [36, 37]. Our results show that the ROH patterns we observe are likely driven by demography (domestication), artificial selection (breed formation), and inbreeding. These processes have resulted in unique ROH patterns within breeds, sharing across breeds, and very little ROH sharing between breed dogs and non-breed village dogs.

We also examine the relationship between ROH and non-disease traits. We observe that the relationship between breed average height and F_ROH_ is positive when using all breed dogs (Fig. 4). This positive effect size opposes recently published work in human populations [55–57]. This may be a result of combining the effect sizes of multiple sizes of dogs (i.e. large and small dogs). When we partition by weight, our results correspond to previous studies on height in breed dogs [58]. As one might expect, we also find a positive β when testing for the relationship between F_ROH_ and weight. We also observe a negative β with respect to the relationship between F_ROH_ and lifespan, indicating an association with inbreeding and survival to old age. This result is consistent with the findings in dogs where an association with disease phenotypes was observed [37]. Thus, when we partition dogs by weight [59], we find a trend where large dog breeds have a shorter lifespan (*β* = -0.15) and small dogs have a longer lifespan (*β* = 0.11). However, our results were not significant (Table S3). Previous studies have linked size with longevity, and this remains an interesting area for future work [60]. When we include lifespan as a covariate when associating F_ROH_ and breed average height in all individuals, we still find a nominally significant, but larger p-value, of 0.034. Further, the effect size of F_ROH_ on height decreases from *β* ∼ 0.31 to *β* ∼ 0.19. We also observe a negative (*β* ∼ -0.83) and significant relationship between height and lifespan (p = 6.41 × 10^-41^).

Our GWAS using ROH identifies several significant associations between SNPs and the quantitative traits: height, weight, and lifespan (Table S6, Table S7, and Table S8). When utilizing data for all individuals, there were three hits *CERS3* (chromosome 3), *CBFA2T2* (chromosome 24) and *SNTA1* (chromosome 24) above the GWS threshold for height. In previous studies, these three genes were found to harbor genetic variants associated with human height [61, 62]. Additionally, human genetics studies have found *CERS3* to be linked with Body-Mass Index (BMI) [63]. We see that many of the significant genic hits within our subgroup of small individuals (*PITPNA*, *LRRC28*, *TTC23*, *HGF*, *SLC43A2*, and *VPS53*) are already associated with human height [62, 64]. We also observe a new relationship between *RPH3AL* and height. Though there is limited research on *RPH3AL,* its role in exocytosis and the secretion of growth hormones in humans has been noted [65]. Importantly, the signal within *IGF1R* replicated, and was above the SWS threshold. A mutation in *IGF1R* was linked with height in small dog breeds, like Chihuahuas [58]. Lastly for small individuals, *COMMD6* was shown to be associated with height. *COMMD6* has primarily been studied in relation to immune function, and to our knowledge, no studies to date have reported any associations between this gene and height [66]. In the subgroup of large individuals, the only gene above the SWS threshold was *PRR16* (chromosome 11), which has been previously associated with human height [62].

For weight, across all our samples, our GWAS results pinpoint *PKD2* (chromosome 32) to be the most strongly associated. Mouse models have shown the presence of *PKD2* has been linked to an increase in fat absorption, potentially promoting obesity [67]. Several other genes, with significant associations in our study, have been linked to weight in other species. *SNTA1* was found to be associated with the Live Body Weight (LBW) of goats, which is a factor often used to assess livestock health [68]. Another gene, *CBFA2T2*, was shown to regulate adipogenesis in both humans and mice, which can affect obesity when its expression is altered [69–71]. *TXNRD3* influences adipocyte differentiation through its involvement in the Wnt signaling pathway, a key process in the regulation of body fat and energy storage [72–74]. *CERS3*, also associated with height, has been linked with Body Mass Index (BMI) in humans [63]. Additionally, the genes *SPIDR*, and *ZNF341* have been associated with total body fat and bone mineral content, respectively [75–77]. *CHMP4B* has been linked to the human birth weight, no studies have directly linked it to obesity and/or body mass. [78, 79]. Similarly, *ZXDC* has primarily been investigated in the context of immune system regulation and cancer biology. To our knowledge, this is the first association with weight thus far, which provides an additional avenue for future researchers to explore. In our subgroup of small individuals, we identify similar associations between weight and the genes *SNTA1*, *ZNF341*, and *CBFA2T2*. Additionally, we identify new associations specific to this subgroup with *PITPNA* (also associated with height) and *SLC43A2*. Both of these genes were shown to decrease body weight in knockout mice [80, 81]. In the subgroup of larger individuals, we find two genes to be associated with weight, *PKD2* and *SPIDR*, both of which are associated with body fat.

For lifespan, we highlight several associations of interest. When testing all samples, we find significant SNPs linked to the genes: *TXNRD3*, *DRD3*, *TIGIT*, *SEPT11* and *ZXDC. TXNRD3* has been previously associated with lower survival rates for various types of cancers [82]. However, to our knowledge, no previous studies have found an explicit connection between *TXNRD3* and lifespan. The gene that encodes the D3 dopamine receptor, *DRD3,* is associated with schizophrenia, which has too been linked with decreased life expectancy [83, 84]. We observe significant signals at *TIGIT* and *SEPT11,* which both have been tied to the promotion of tumor growth and reduced longevity [85–87]. *ZXDC* was previously associated with cervical cancer metastasis, which has a significant negative impact on patient survival [88]. In small dogs, we saw an association with *RUNX2*, which has also been associated with multiple cancers and poor patient prognosis [89]. In large individuals, our GWAS revealed *SLC41A3* to be associated with lifespan. A relationship between this gene and liver hepatocellular carcinoma, which is the second primary contributor of carcinoma-associated death, has been established [90]. Another gene found in the group of large individuals is *CFAP100*, which to our knowledge has not previously been linked to lifespan.

Overall, our GWAS suggests that quantitative traits in dogs are polygenic with multiple variants associated with a trait, and some traits are not entirely additive. The newly identified genes, which have not been observed in previous studies, could be used as candidates in future functional exploration. The presence of these genes also highlights new avenues to explore complex trait architecture and non-additivity in domesticated species. In humans, there have been 25,551 associations with height, 2,233 associations with weight, and 664 associations with lifespan as of 2023 [91]. After correcting for inflation, we recover a total of 50 associations with height, 84 associations with weight, and 21 associations with lifespan. This corresponds to previous work, which has suggested that dogs have a more simplified complex trait architecture than humans. [38, 92].

Importantly, we conducted an additional test to validate our ROH based GWAS by attempting to replicate a previously identified peak that was known to be homozygous on chromosome 13 within RSPO2 for furnishings in dogs [29]. We can identify the same peak on chromosome 13 for RSPO2 (Fig. S26), though it did not meet genome-wide or suggestive significance after p-values were corrected (Fig. S27). In addition to this peak, we observe additional peaks that had not been previously identified. Replicating this peak suggests our approach, which corresponds to different information, is powerful and worthwhile. Notably, similar power in human GWAS requires hundreds of thousands or even a million individuals [61, 62]. Here, we capture non-additivity when using ROH-mapping GWAS, and we can capture this information with less than 1000 dogs. This is due to the unique domestication and breed formation bottlenecks that dogs experienced that simplified complex trait architecture [29, 38, 43, 93–95].

There are some limitations in the work that we would like to highlight. First, the phenotypic values used in the analysis were calculated based on the breed average, restricting us to study associations between breeds. In other words, the within breed values would not yield any significant results as the expected association between the trait and F_ROH_ would be 0, due to using a breed average phenotype. Using the breed average has been previously validated [29, 38, 43, 93–95] but the assumption does come with caveats. For example, when using a breed average value, we are ignoring variance among individuals within a breed. This variance within the breed is quite small due to the strict standards of the AKC. To this end, we mathematically show that if the within breed variance is small, the effect size computed from individual level data will be equivalent to the effect size from using the average. However, bias can be introduced in some cases when aggregating across breeds. Secondly, breed group categorization based on both neighbor-joining trees and AKC could potentially still introduce errors when grouping some breeds. Additionally, we did observe that there was a concentration of peaks in the last few chromosomes (Fig. S8, Table S9). We believe that the large number of peaks on the smaller chromosomes could be due to stronger linkage disequilibrium in the smaller chromosomes, thus amplifying the strength of the signal. Alternatively, some chromosomes (such as chromosome 32) have a higher GC content when examining the CanFam3.1 assembly [96], which could introduce noise into our results. Despite these limitations, the significant relationship between F_ROH_ and certain phenotypic traits suggest that inbreeding from domestication and artificial selection played a strong role in shaping complex non-disease traits within breed dogs.

In summary, our work has further elucidated the genetic structure of quantitative trait architecture in dogs. We also highlight how inbreeding, quantified in the form of ROH, occurred alongside selection for extreme phenotypes during breed formation. These ROH tag regions of the genome associated with complex non-disease phenotypes.

## Methods

### Data filtering and categorizations

In this study, we utilized whole genome sequences from 722 canids, which included various wild species, dingoes, and domestic dogs [29]. This data can be accessed via NCBI accession number PRJNA448733. All filtering was accomplished using BCFtools [97] where we retained only biallelic SNPs and sites of genotype quality score greater or equal to 20. Additionally, we removed any missing genotype rates above 10% and removed variants from sex chromosomes. Our filtered data consisted of 4,053,761 biallelic loci for all 38 pairs of autosomal chromosomes. Given the nature of our study, we removed all wild canids, mixed-breed individuals, and samples for which breed information was unavailable. Each remaining sample was categorized by common name or breed [29]. These classifications were informed by neighbor joining trees of domestic dogs [21] and American Kennel Club (AKC) groupings [98] . This resulted in 13 breed groups namely, Ancient Spitz (31 individuals and 11 breeds), Herding (76 individuals and 15 breeds), Mastiff-like (65 individuals and 15 breeds), Retrievers (50 individuals and 6 breeds), Scent Hound (23 individuals and 10 breeds), Small Terrier (102 individuals and 8 breeds), Terriers (16 individuals and 7 breeds), Spaniels (22 individuals and 8 breeds), Toy Dogs (18 individuals and 7 breeds), Working Dogs (44 individuals and 10 breeds), Sight Hound (21 individuals and 9 breeds), Village Dogs (75 individuals and 13 regions), and Chinese Indigenous dogs (15 individuals). Overall, our working dataset included 558 individuals and 106 different breeds (Table S10).

### Calling Runs of Homozygosity

To call runs of homozygosity (ROH) we used a likelihood-based inference method called GARLIC v1.6.0a [99]. This method uses a logarithm of odds (LOD) score measure of autozygosity, which is applied in a sliding window for the entire genome [100]. GARLIC requires input data with genotype and sample information in the form of TPED and TFAM files, which were generated using PLINK 2.0 [101] We generated a TGLS file to obtain per-genotype likelihoods. We employed genotype likelihood data in the form of GQ to account for errors in phred-scaled probability. Next, a window size of 100 SNPs was chosen based on SNP density (Fig. S28), with the window incrementally advancing by 10 SNPs at each step. GARLIC offers built-in ROH length classification, for which we defined 6 classes: <1 Mbp, 1-2 Mbp, 2-3 Mbp, 3-4 Mbp, 4-5 Mbp, and >5 Mbp. To account for sample size differences in allele frequency estimation, we set the number of resamples to 40. All other flags were set to default, and the following parameters were used: --auto-winsize --auto-overlap-frac --winsize 100 --centromere --size-bounds 1000000 2000000 3000000 4000000 5000000 --tped --tfam --tgls –gl-type GQ -- resample 40 --out. To eliminate segments that were noisy, very short, and common, we filtered to retain ROH segments longer than 0.5 Mbp.

We conducted independent ROH calling for breeds with at least 10 individuals. Breeds consisting of fewer than 10 members were integrated into their respective breed groups (Table S11). Subsequently, ROH calls from all breeds, irrespective of individual numbers, were combined into their respective breed groups for further analysis. For each individual, we examined: 1) the relationship between coverage and F_ROH_ (Fig. S29) and 2) the relationship between the total number of runs of homozygosity (n_ROH_) and total length of runs of homozygosity (s_ROH_) (Fig. S30)).For each individual and breed group, we calculated the arithmetic mean and range of both total n_ROH_ and total s_ROH_, while binning them based on ROH size classification. The longest total s_ROH_ (2141.22 Mbp) and the highest total n_ROH_ (1039) were identified in two individuals - PER00747 and PER00393, respectively - from the small terrier Yorkshire breed. PER00747 was sequenced at a low depth (∼2x), which inflated s_ROH_. To mitigate the effects of low depth PER00747 and PER00393 were removed from downstream analysis.

### Inbreeding coefficient

For each sample and breed group, we computed the inbreeding coefficient, F_ROH_, defined as the fraction of autosomal genome in ROH regions [102]:

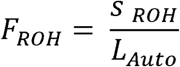

Where s_ROH_ is the sum of the length of runs of homozygosity in an individual’s genome and L_Auto_ is the length of autosomal genome in base pairs. An autosomal genome length of 2,203,764,842 base pairs was used since variants were called with the CanFam3.1 (NCBI RefSeq assembly: GCF_000002285.3) reference genome.

### ROH Sharing Matrix

We generated an ROH sharing matrix, a square matrix that quantifies the sum of ROH overlaps between two individuals. This allowed us to analyze shared ROH patterns among dog breeds. To populate this matrix, we first generated individual ROH bed files for each sample. We then used BEDTools [103] to identify overlapping regions between pairs of ROH data. The following parameters were used: bedtools intersect -wao.

### Phenotype data

We updated the phenotypic data [29] for 13 traits: bulky, drop ears, furnish, hairless, height, large ears, length of fur, lifespan, long legs, muscled, weight, white chest, and white head. For the three continuous traits, namely height, weight, and lifespan, we used average values obtained from AKC [98] (Table S12). We chose to use breed average values because this was previously shown to be a reliable measure for association tests [29, 38, 43, 93–95]. When sex-specific information was available, it was incorporated into the dataset; otherwise, we applied the same values for both males and females. We applied min-max scaling to normalize the three continuous traits and further categorized them into small and large groups based on the average weight (21.66kg) of all individuals (Tables S13-S15). For the remaining 10 traits, we averaged the values by the phenotypic data available for their respective breeds (Table S16). We used binary encoding to assign a value of 1 to individuals expressing the trait and 0 to those who did not.

### Association tests

We computed the association between F_ROH_ and the 13 phenotypes using GMMAT, an R package that performs association tests with generalized linear mixed models (GLMM) [104]. To fit the GLMM, we used the built-in function glmmkin, which allowed us to examine the continuous traits (height, weight, and lifespan) as the quantitative phenotype traits. This also enabled us to include individual F_ROH_ as a covariate and use the ROH sharing matrix as a kinship matrix. We fit the model assuming a Gaussian distribution for the continuous phenotypes and used the identity link function. For associations between height, weight, and lifespan with F_ROH_, we used the updated average phenotypic values of the breed for all 466 individuals (Fig. S31 and Supplementary Text). Based on our association tests and previous studies [105, 106], weight and F_ROH_ have been observed to have a significant association, thus we included them as covariates in our models. Weight and the interaction of weight and F_ROH_ were used as covariates across three data subsets (all individuals, small individuals, and large individuals) (Table S4). To fit the GLMM to the 10 binary traits, we specified the binomial distribution as the family and used the logit link function. Samples with no information available on the presence or absence of these phenotypes were removed from analysis. Due to lack of phenotype information, Chinese indigenous dogs and village dogs were excluded from the association tests.

### ROH-mapping GWAS

To explore the biological mechanisms through which ROH-associated SNPs influence phenotypic traits, we conducted Genome-Wide Association Studies (GWAS). We generated a dataset that examined the occurrence of SNPs in ROH regions using BCFtools intersect and BCFtools subtract [97]. Any SNPs located outside of ROH regions were removed. Using GMMAT, we fit GLMM models assuming a recessive genetic model for quantitative traits (height, weight, and lifespan), incorporating ROH-associated SNPs as covariates, an ROH sharing matrix for kinship structure, and a Gaussian distribution with an identity link function [104]. Additionally, given that breed structure has a particularly strong effect in dogs, we utilized a kinship matrix quantified by pairwise ROH sharing between individuals when associating traits with F_ROH_. This procedure follows from an approach from a study on disease traits [37]. Lastly, we fit GLMM models using weight and the product of weight and F_ROH_ as covariates for the three data subsets (all individuals, small individuals, and large individuals) (Table S5). To obtain the effective sample size, adjusted for auto correlations between sites, we used the option effectiveSize within the coda R package [107]. To calculate the number of independent tests we used p-values for every SNP as the input, thus incorporating autocorrelation caused by LD. This approach was shown to be effective for correcting for structure previously [108]. To obtain the Genome-Wide Significance (GWS) and Suggestive-Wide Significance (SWS) thresholds, we used the Bonferroni correction with respect to effective sample size [109–111]:

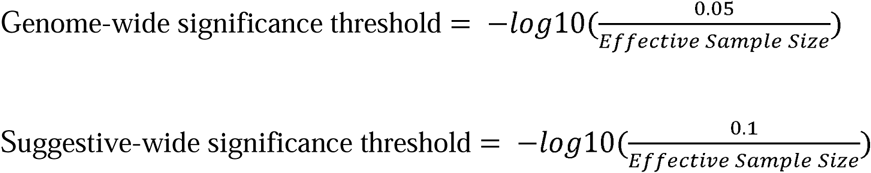

To correct for population stratification, we calculated the genomic inflation factor, λ [112, 113]. We corrected the log-scaled p-values by dividing them by the genomic inflation factor (Table S5). To visualize the GWAS summary statistics, we used an R package, qqman to create Q-Q and Manhattan plots [114].

## Accession Numbers

Whole genome sequence data is available on NCBI, accession number: PRJNA448733. The source phenotype data was obtained from [29]. All summary data are contained within the article and its supplementary information.

## Declarations

The authors have nothing to declare.

## Acknowledgements

Sweetalana would like to thank the entire Szpiech Lab for their invaluable support.

## Availability of Data and Materials

All code is available at https://github.com/jaam92/QuantTraitInDogs and all data has been published.

## Author Contributions

ZAS and JAM designed and supervised the research project. S performed the research and analyzed data. SN reviewed domestication literature for writing and editing of the manuscript. SH generated the proof and plots for the effect of using breed averages. S along with the assistance of SN, ZAS and JAM wrote the paper.

