## SupplementaryFigures for "Genotypic and phenotypic consequences of domestication in dogs"

### Supplementary Figures

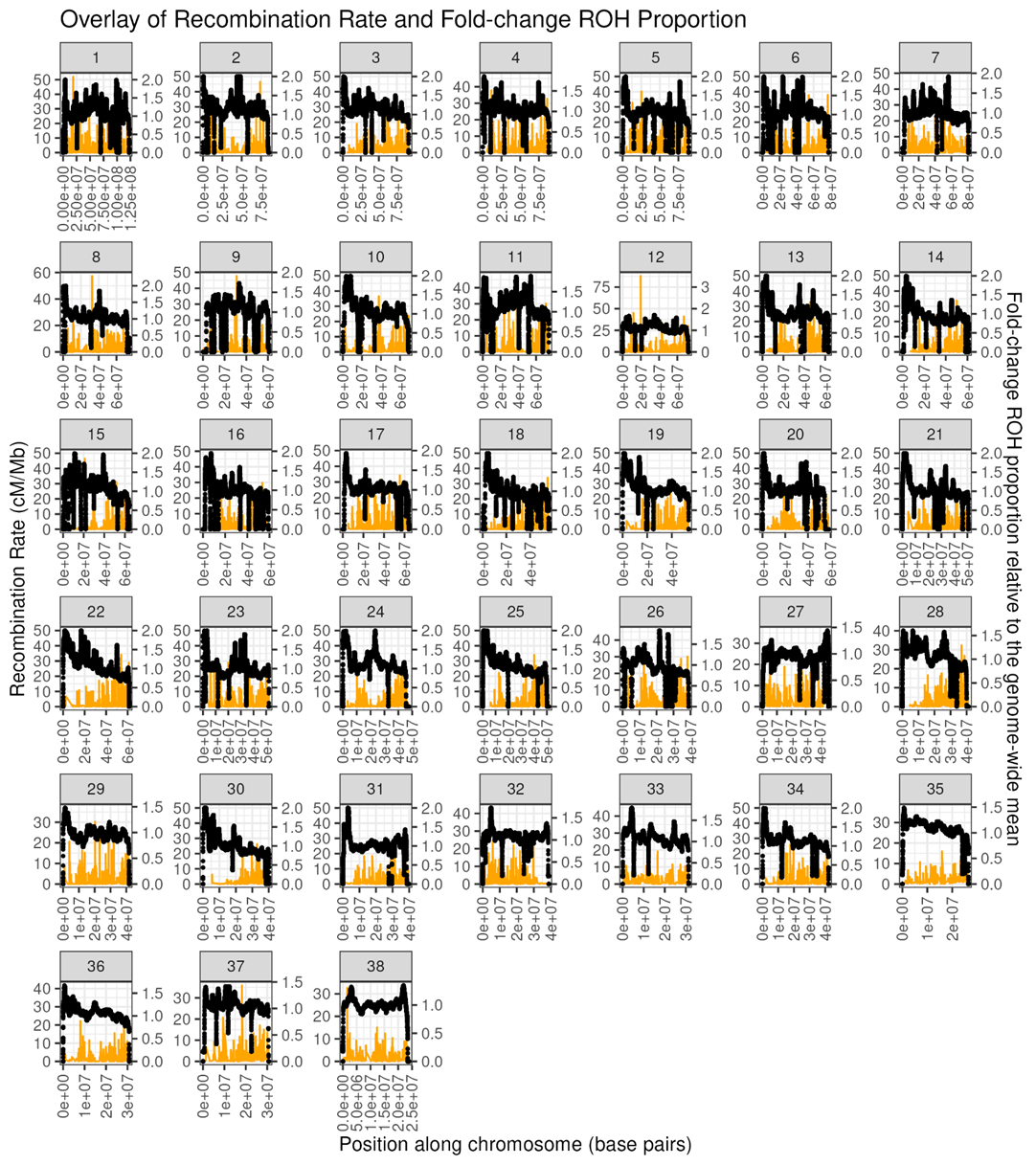

Figure S1. Overlap of recombination rate and ROH fold-change relative to the genome wide average for all samples. The x-axis represents the position along the chromosome in base pairs. The y-axis represents the recombination rate in cM/Mb. Each clade represents an autosomal chromosome.

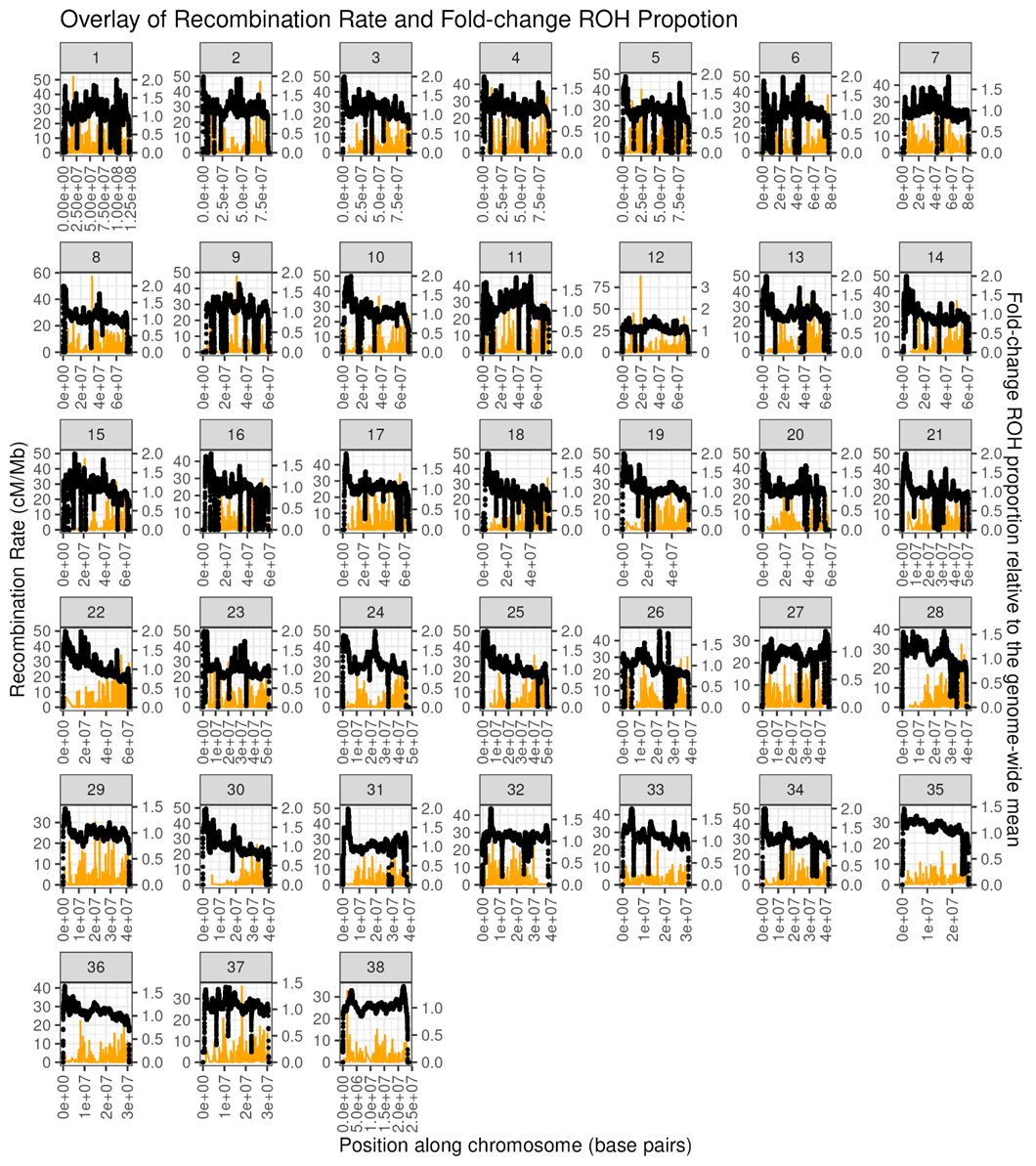

Figure S2. Overlap of recombination rate and ROH fold-change relative to the genome wide average for breed dog only (exclude Chinese indigenous and village dogs). The x-axis represents the position along the chromosome in base pairs. The y-axis represents the recombination rate in cM/Mb. Each clade represents an autosomal chromosome.

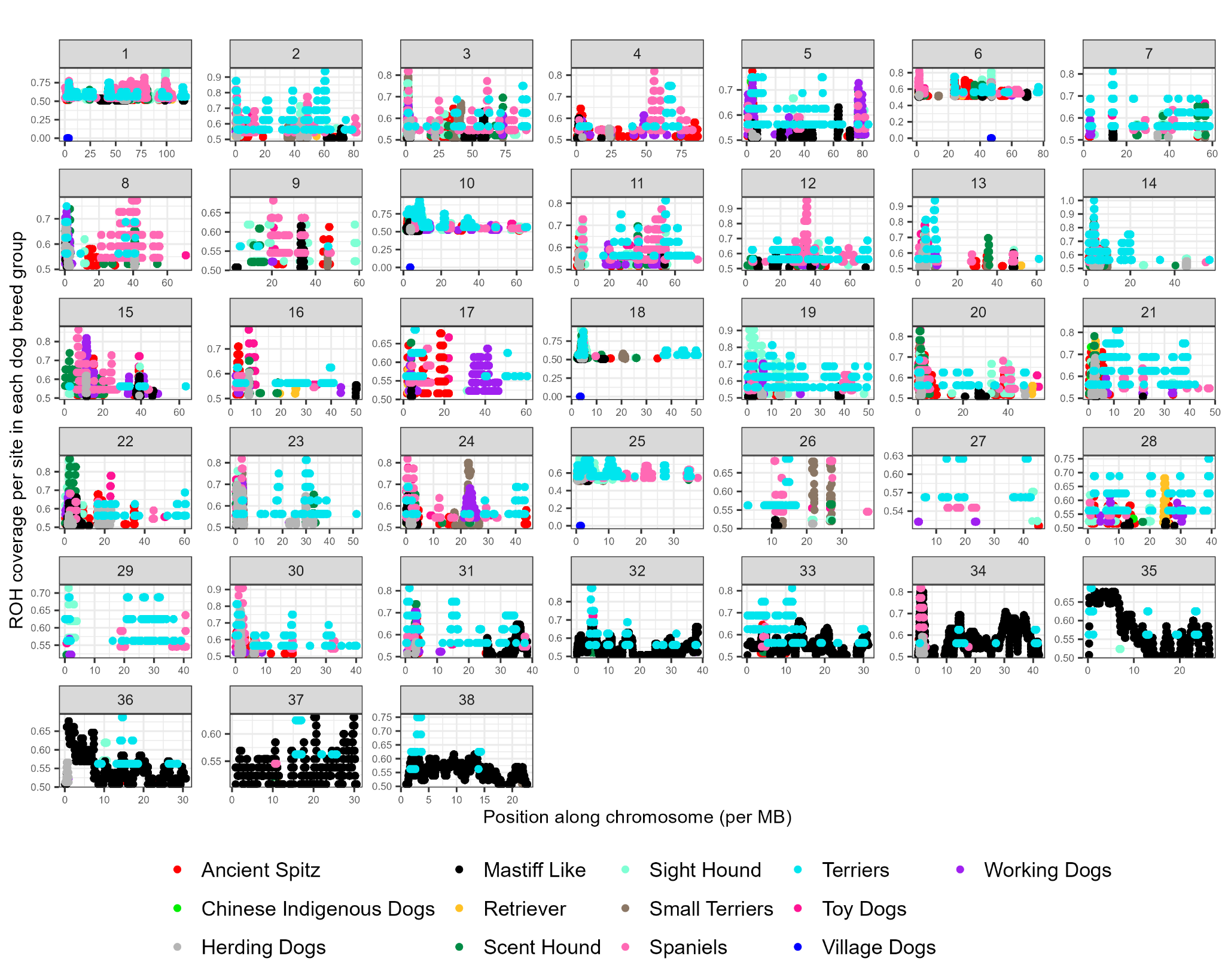

Figure S3 ROH coverage per site for each dog breed group. The y-axis represents the ROH coverage per site (>0.5) for each chromosome. The x-axis represents the position along the chromosome in Mb. Each facet represents an autosomal chromosome.

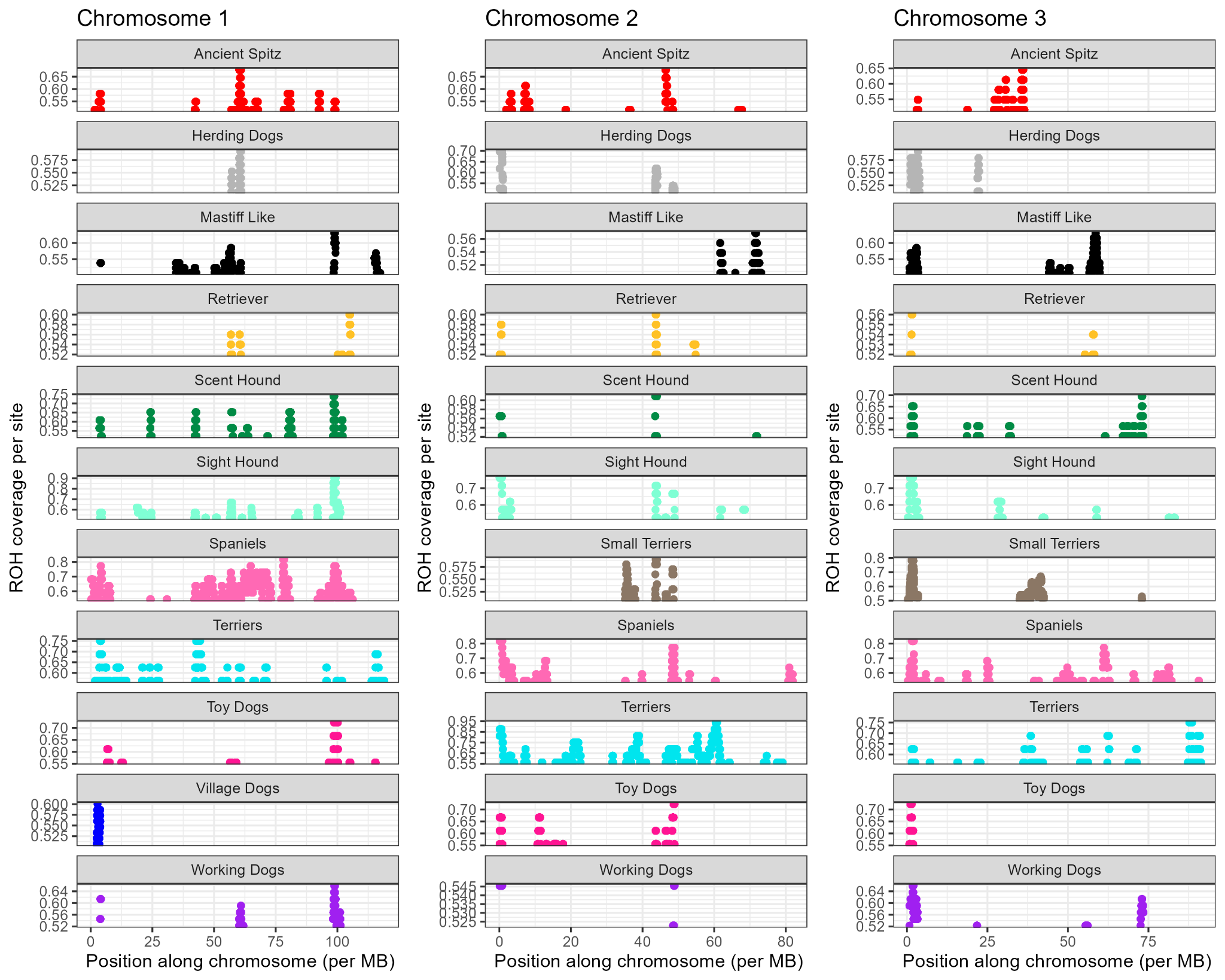

Figure S4. ROH coverage per site for each dog breed group for chromosomes 1 to 3. The y-axis represents the ROH coverage per site (>0.5) for each chromosome. The x-axis represents the position along the chromosome in Mb.

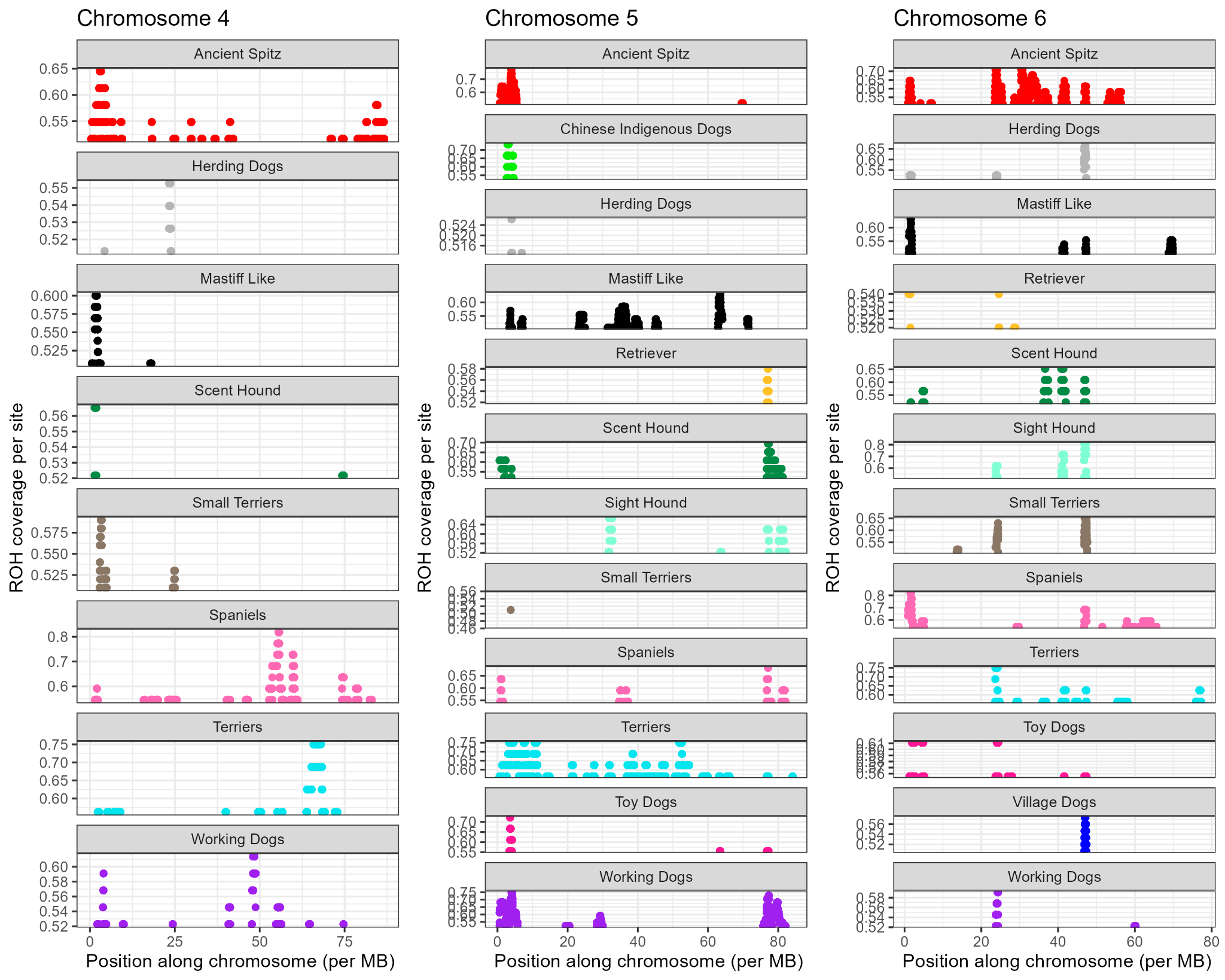

Figure S5. ROH coverage per site for each dog breed group for chromosomes 4 to 6. The y-axis represents the ROH coverage per site (>0.5) for each chromosome. The x-axis represents the position along the chromosome in Mb.

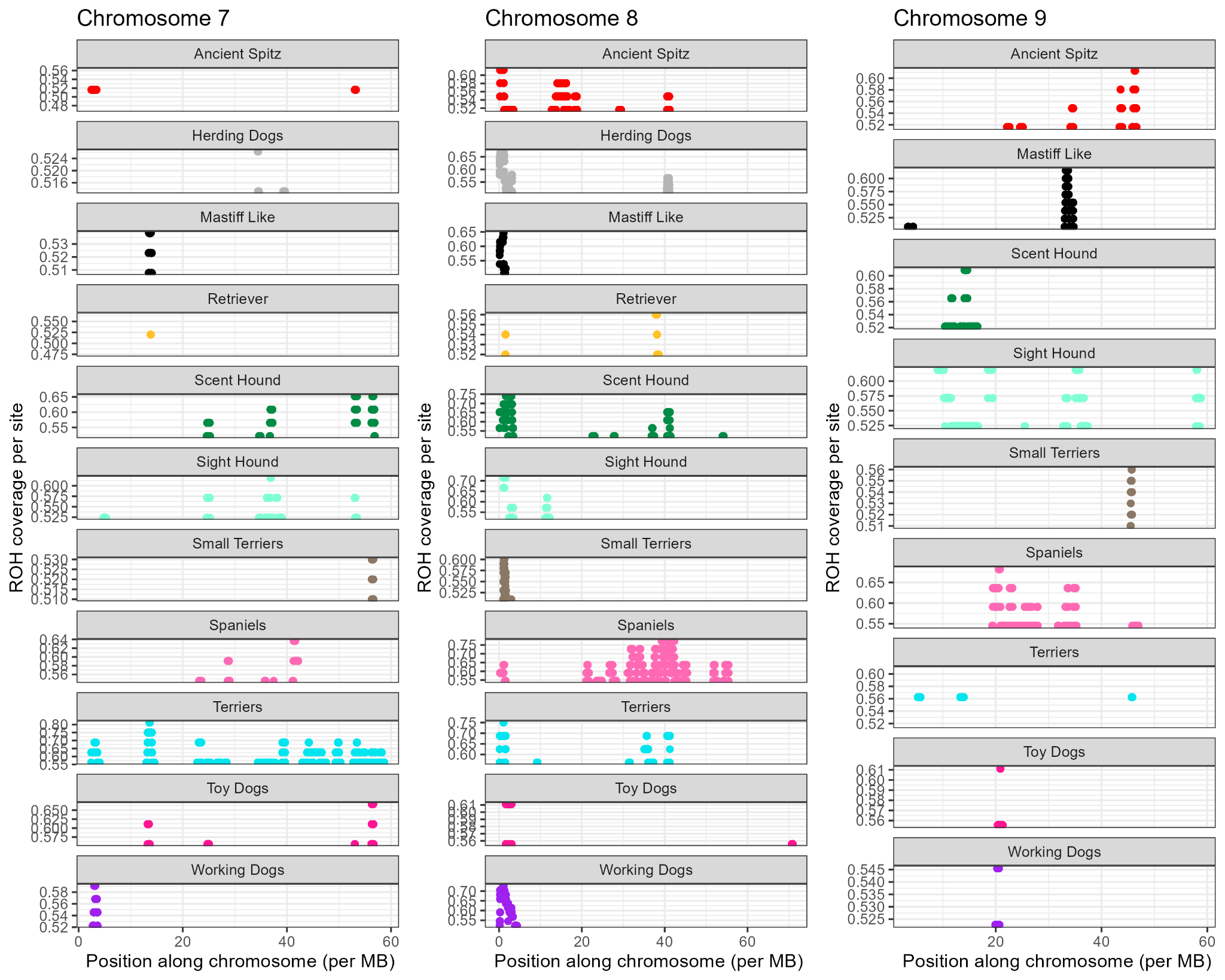

Figure S6. ROH coverage per site for each dog breed group for chromosomes 7 to 9. The y-axis represents the ROH coverage per site (>0.5) for each chromosome. The x-axis represents the position along the chromosome in Mb.

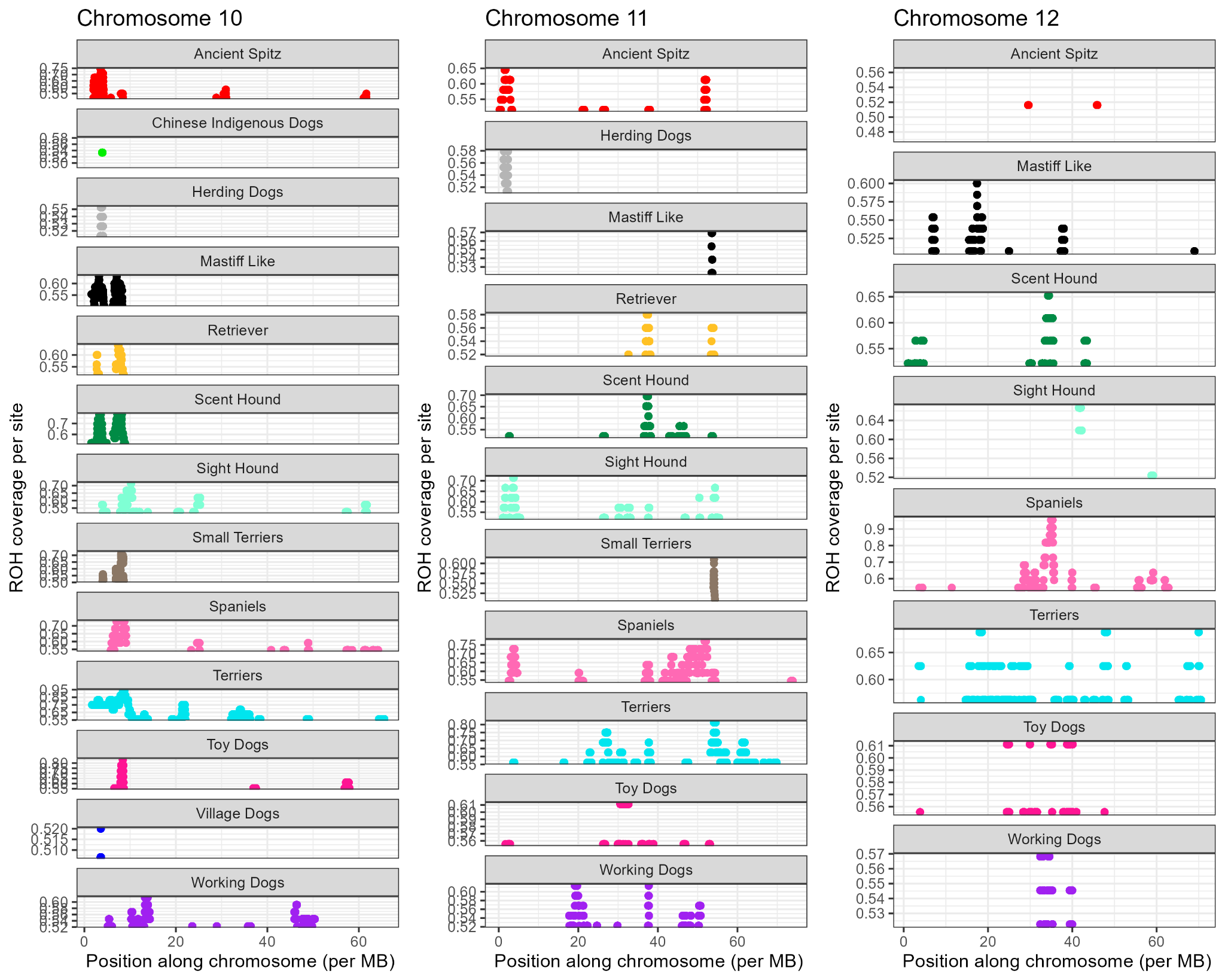

Figure S7. ROH coverage per site for each dog breed group for chromosomes 10 to 12. The y-axis represents the ROH coverage per site (>0.5) for each chromosome. The x-axis represents the position along the chromosome in Mb.

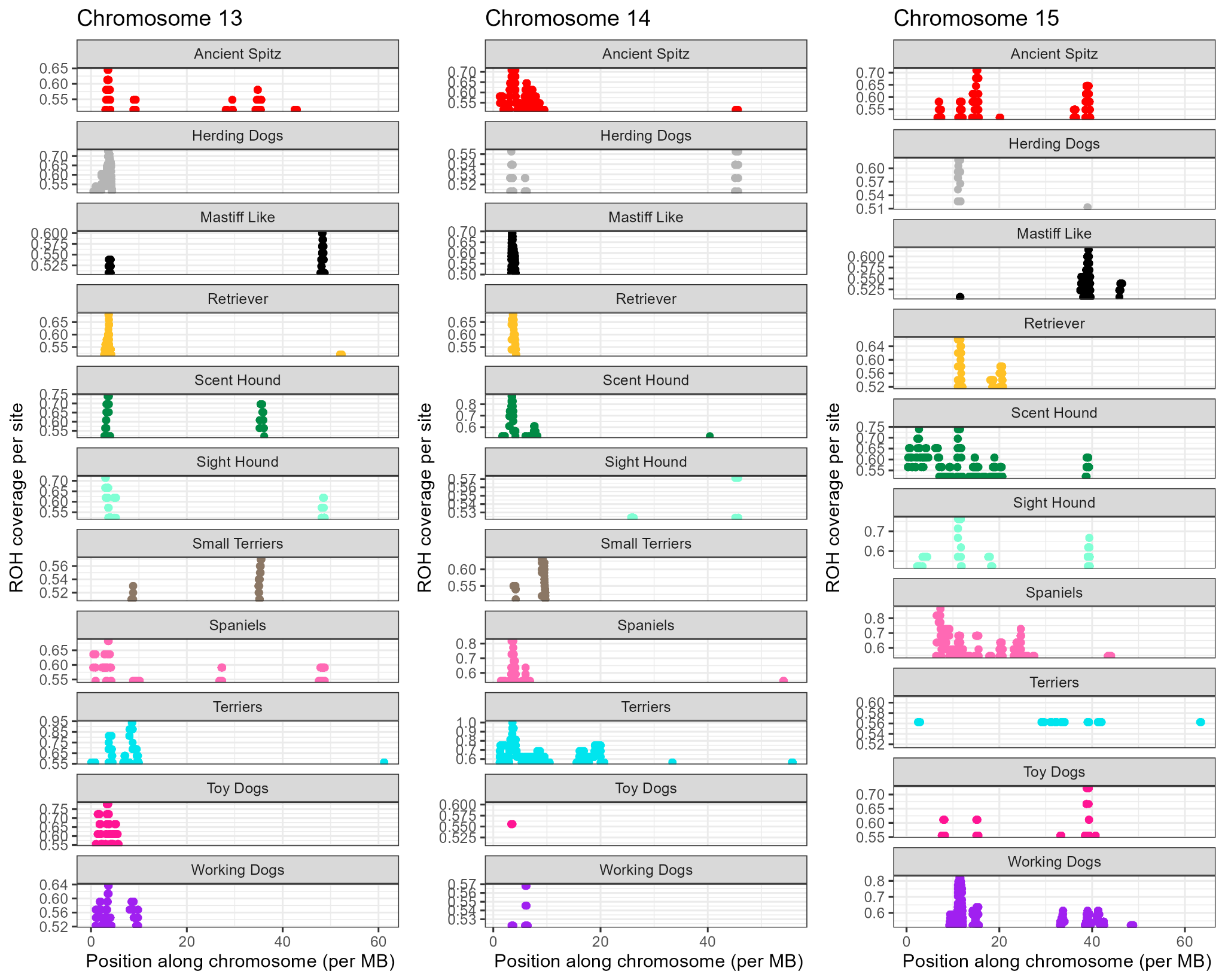

Figure S8. ROH coverage per site for each dog breed group for chromosomes 13 to 15. The y-axis represents the ROH coverage per site (>0.5) for each chromosome. The x-axis represents the position along the chromosome in Mb.

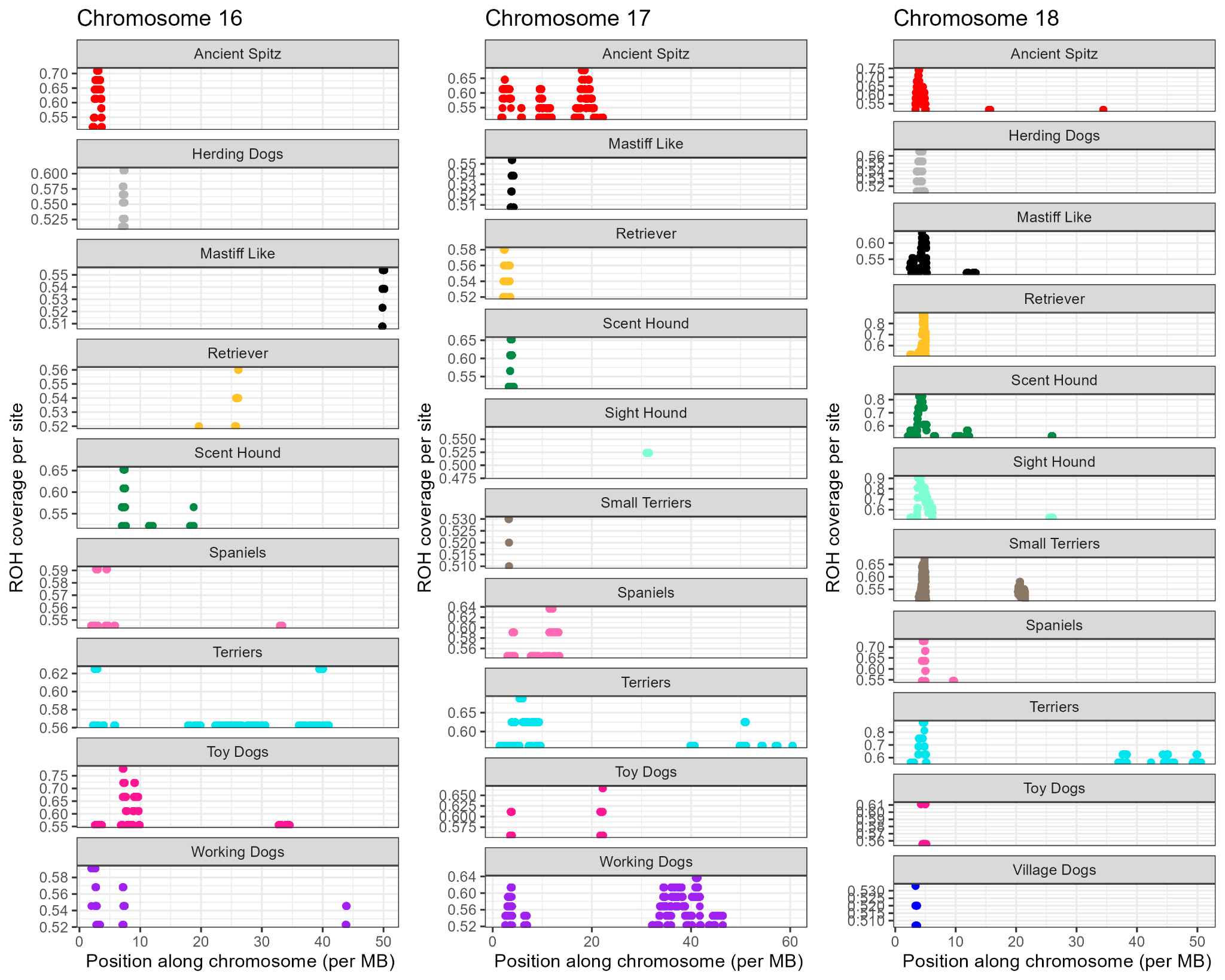

Figure S9. ROH coverage per site for each dog breed group for chromosomes 16 to 18. The y-axis represents the ROH coverage per site (>0.5) for each chromosome. The x-axis represents the position along the chromosome in Mb.

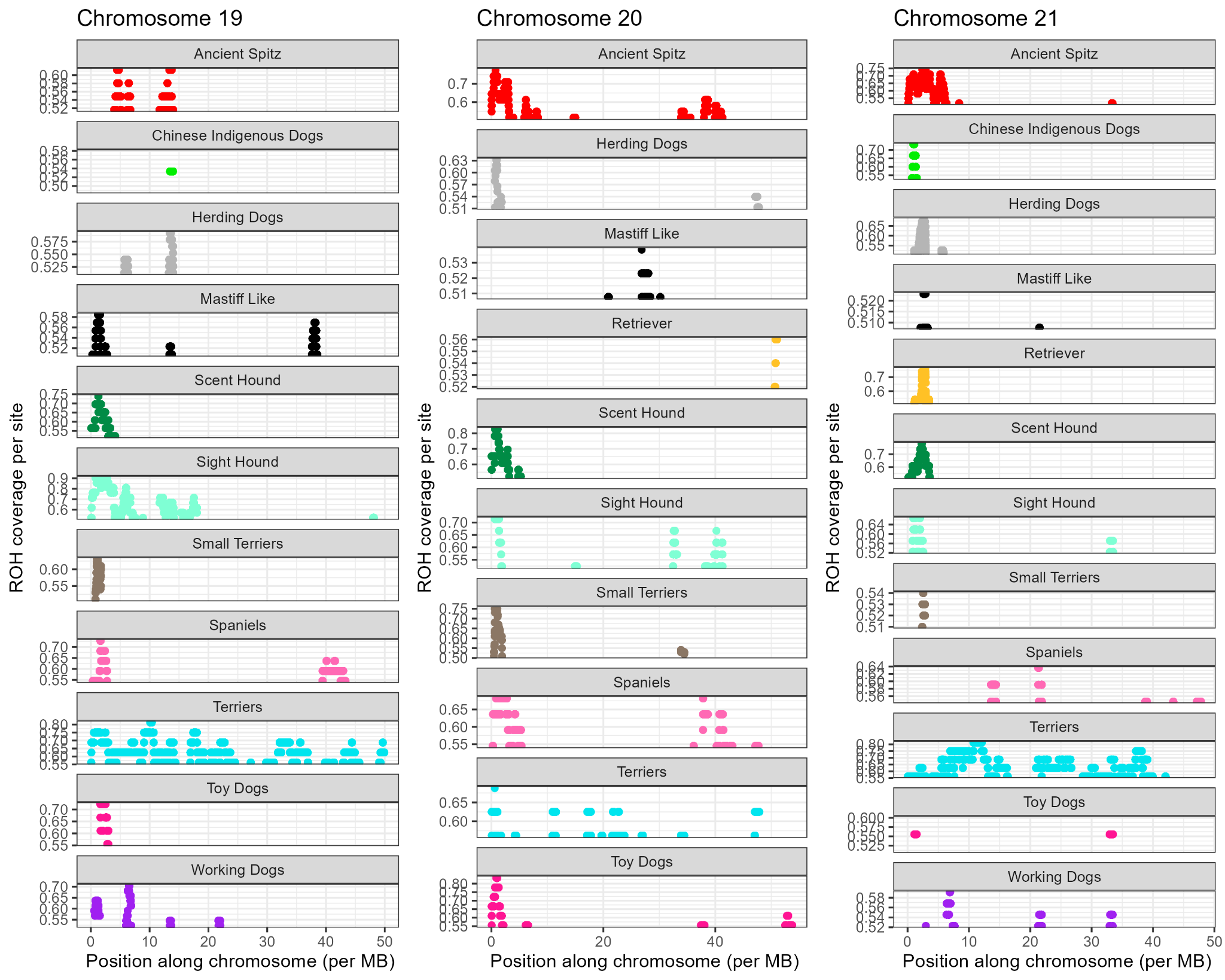

Figure S10. ROH coverage per site for each dog breed group for chromosomes 19 to 21. The y-axis represents the ROH coverage per site (>0.5) for each chromosome. The x-axis represents the position along the chromosome in Mb.

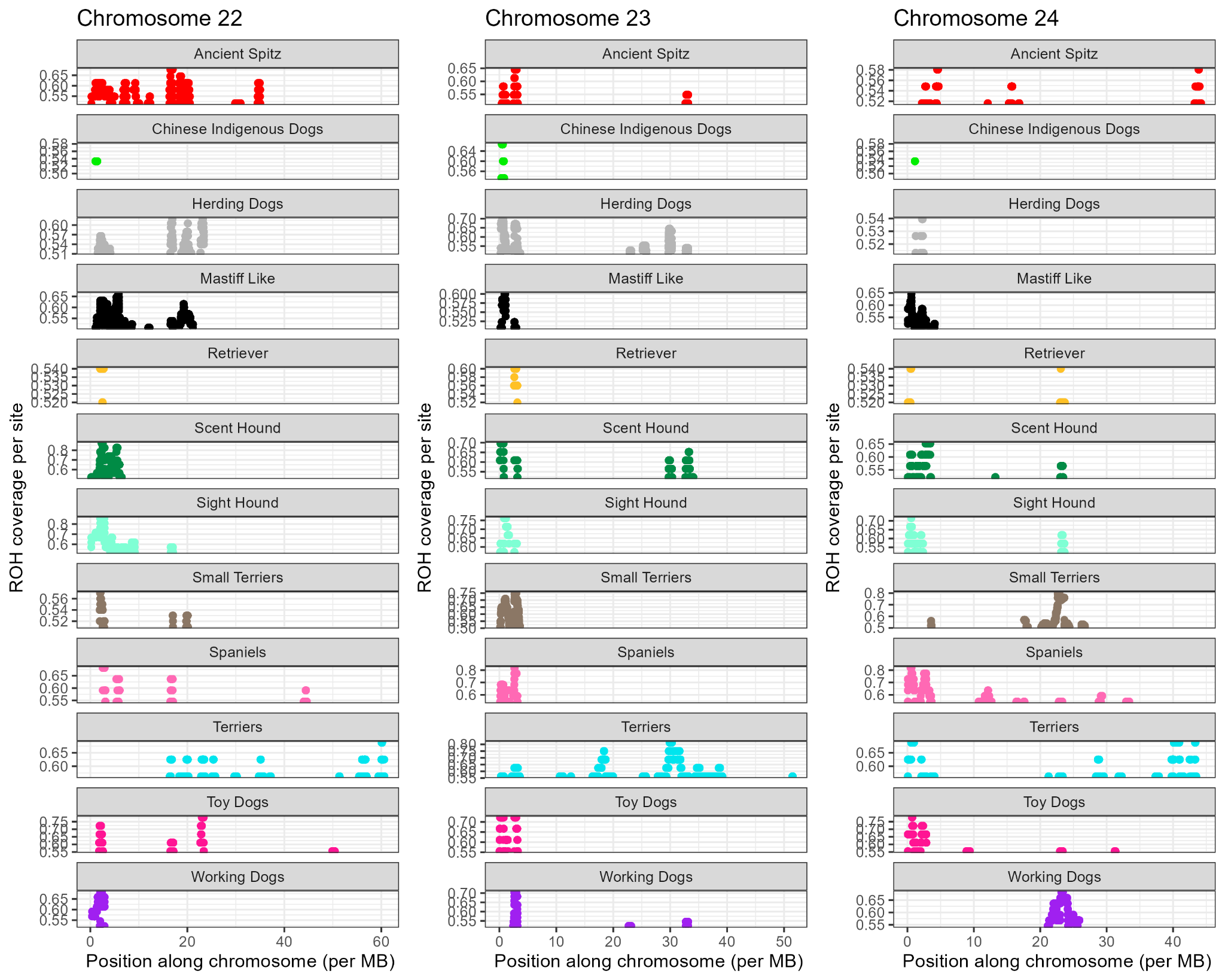

Figure S11. ROH coverage per site for each dog breed group for chromosomes 22 to 24. The y-axis represents the ROH coverage per site (>0.5) for each chromosome. The x-axis represents the position along the chromosome in Mb.

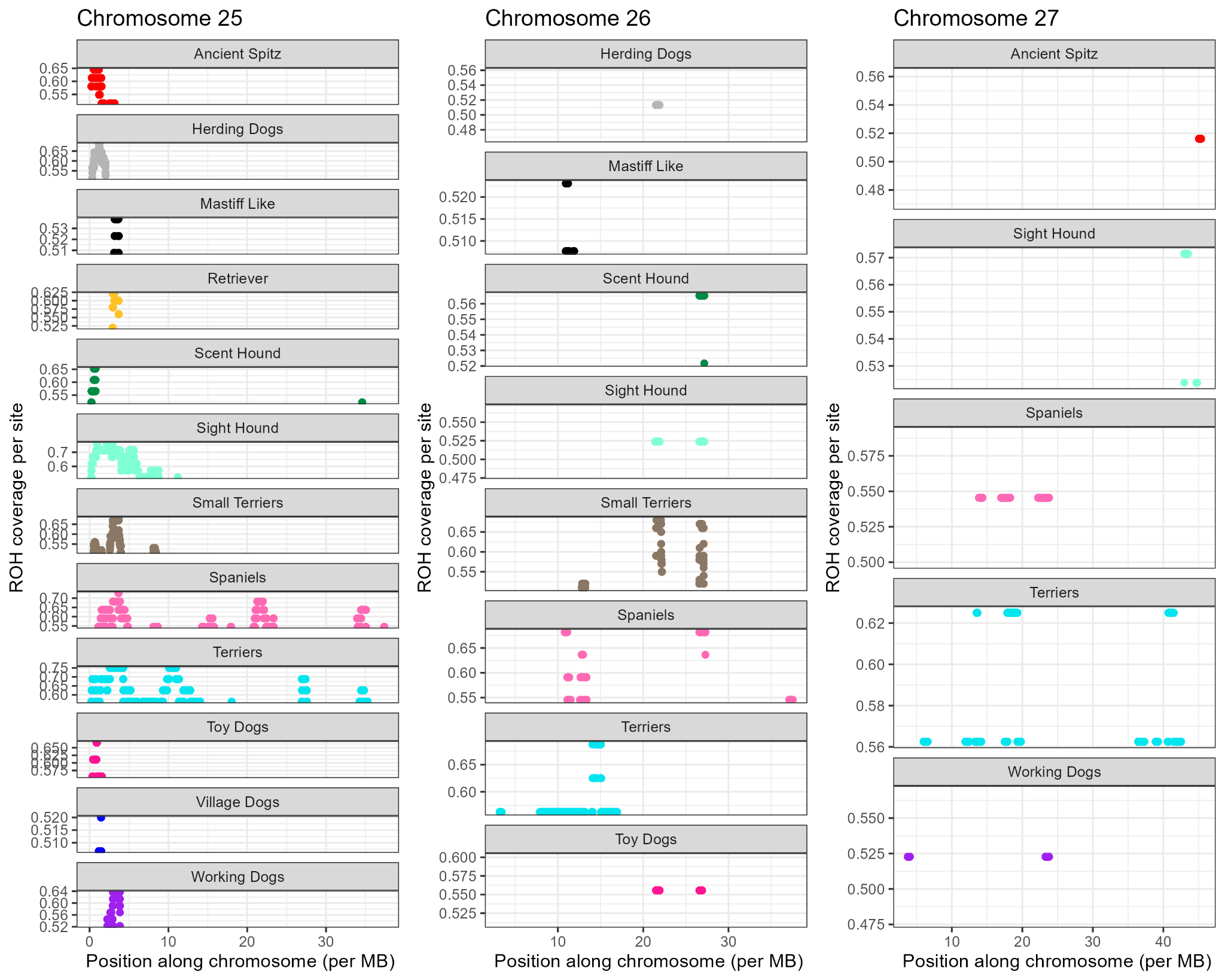

Figure S12. ROH coverage per site for each dog breed group for chromosomes 25 to 27. The y-axis represents the ROH coverage per site for each chromosome. The x-axis represents the position along the chromosome in Mb.

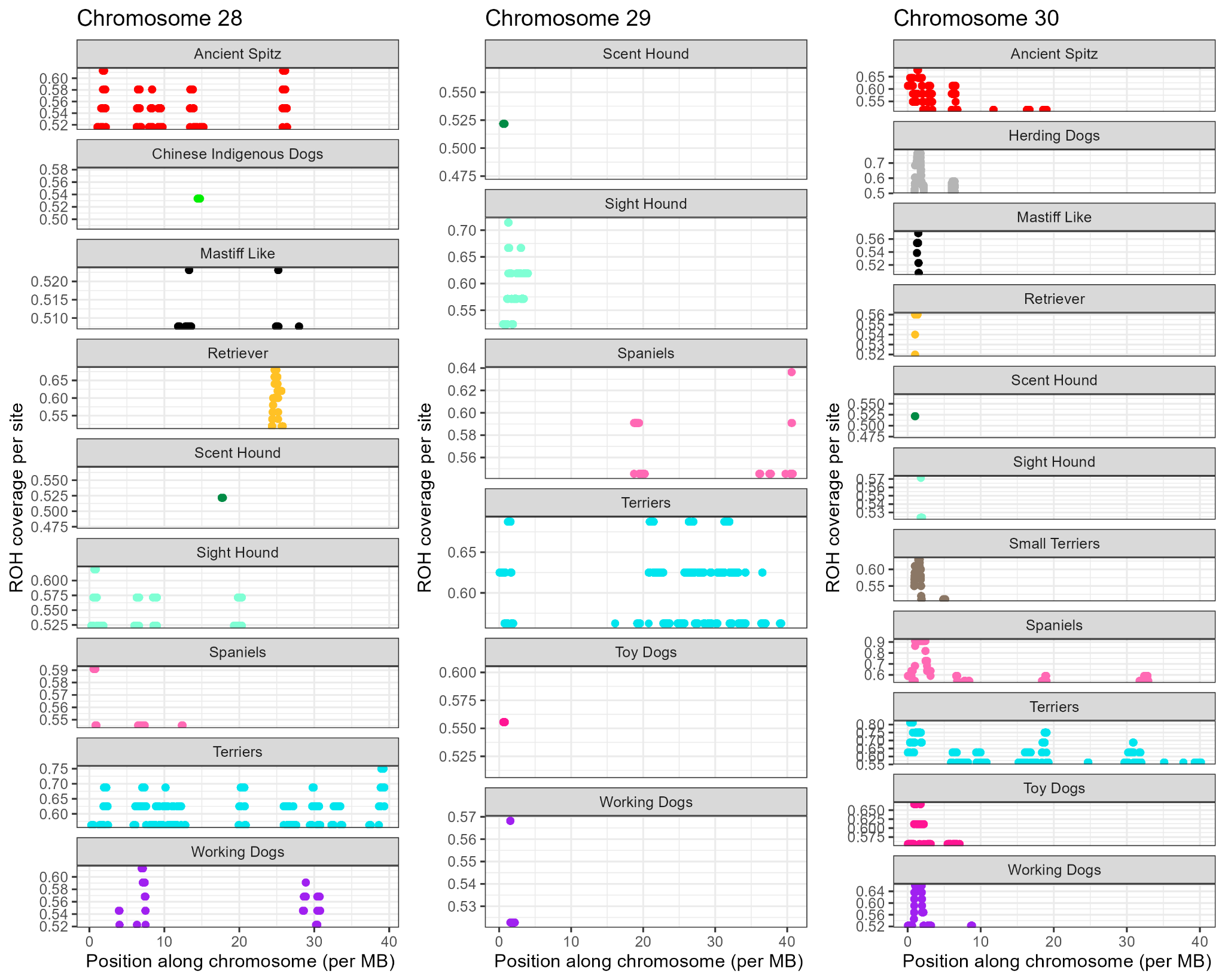

Figure S13. ROH coverage per site for each dog breed group for chromosomes 28 to 30. The y-axis represents the ROH coverage per site for each chromosome. The x-axis represents the position along the chromosome in Mb.

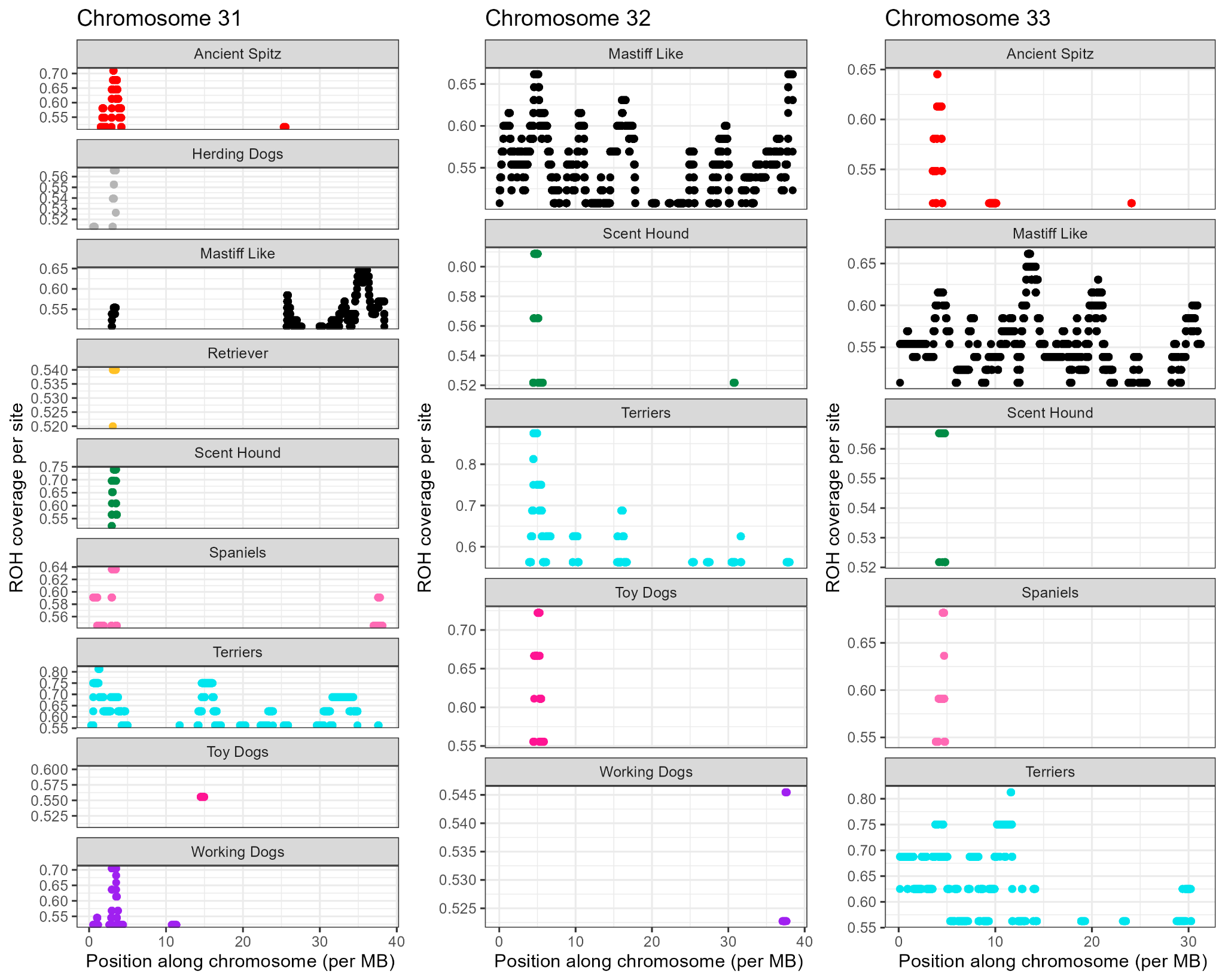

Figure S14. ROH coverage per site for each dog breed group for chromosomes 31 to 33. The y-axis represents the ROH coverage per site for each chromosome. The x-axis represents the position along the chromosome in Mb.

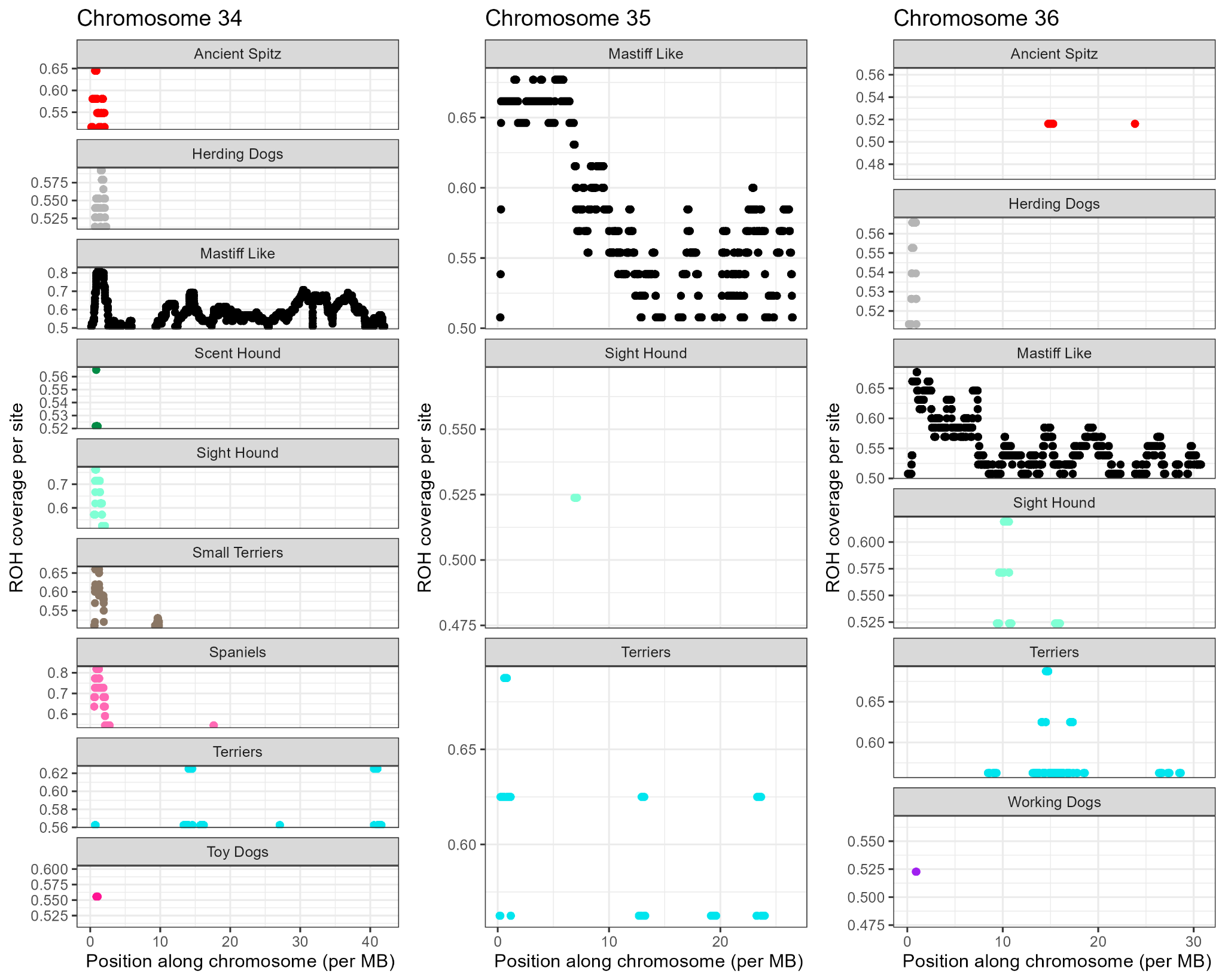

Figure S15. ROH coverage per site for each dog breed group for chromosomes 34 to 36. The y-axis represents the ROH coverage per site for each chromosome. The x-axis represents the position along the chromosome in Mb.

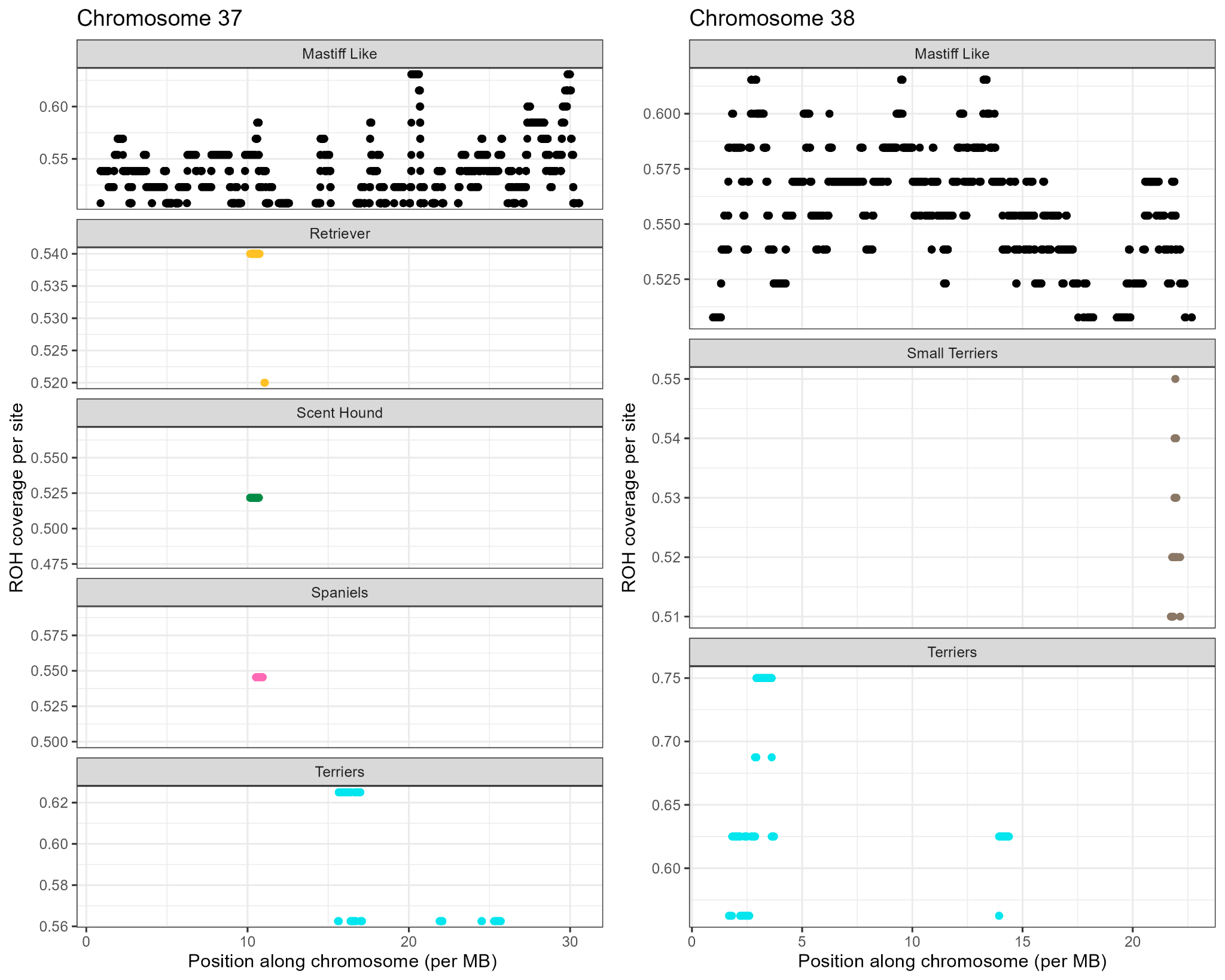

Figure S16. ROH coverage per site for each dog breed group for chromosomes 37 and 38. The y-axis represents the ROH coverage per site for each chromosome. The x-axis represents the position along the chromosome in Mb.

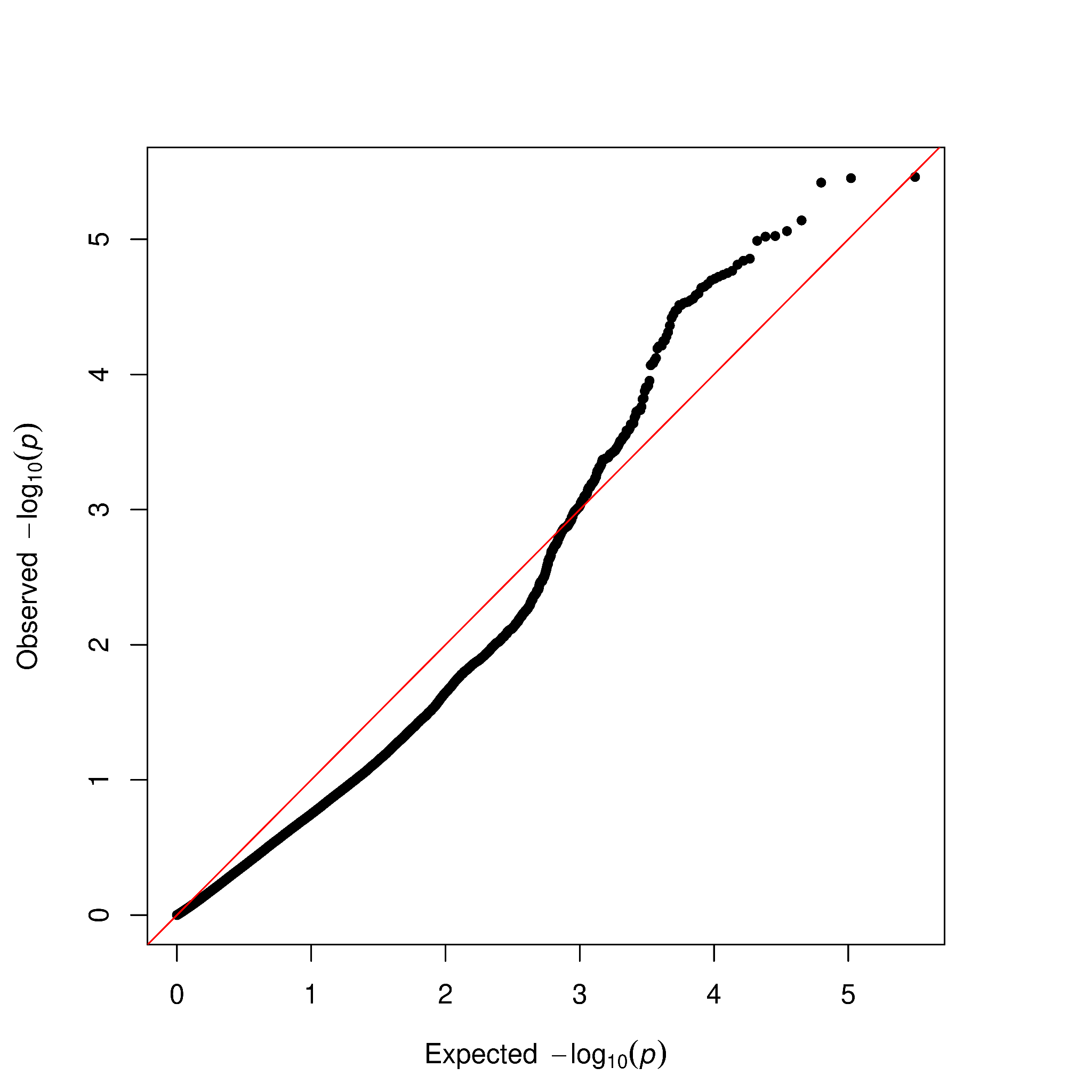

Figure S17. Q-Q plot for GLMM-based GWAS for body height (all individuals).

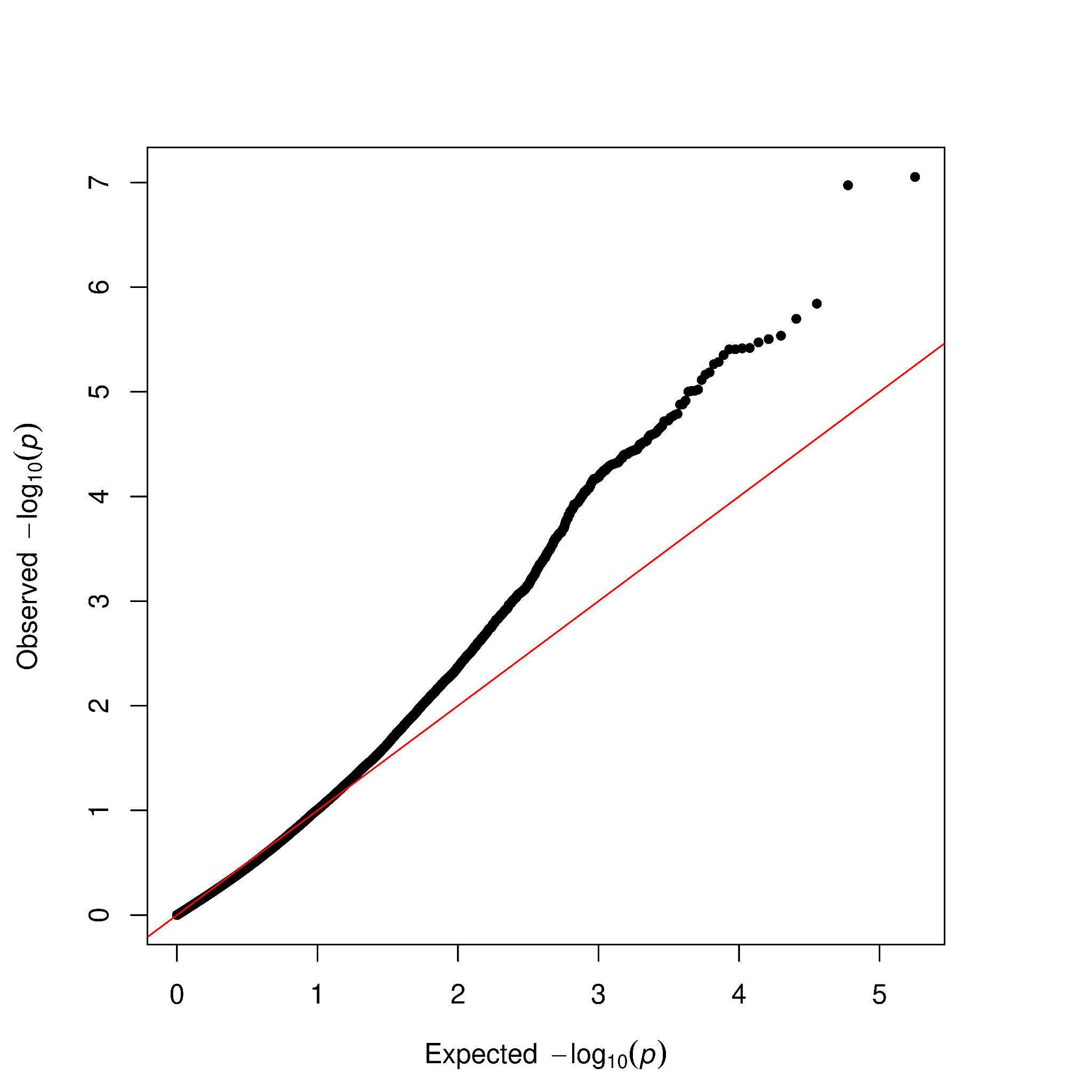

Figure S18. Q-Q plot for GLMM-based GWAS for body height (small individuals).

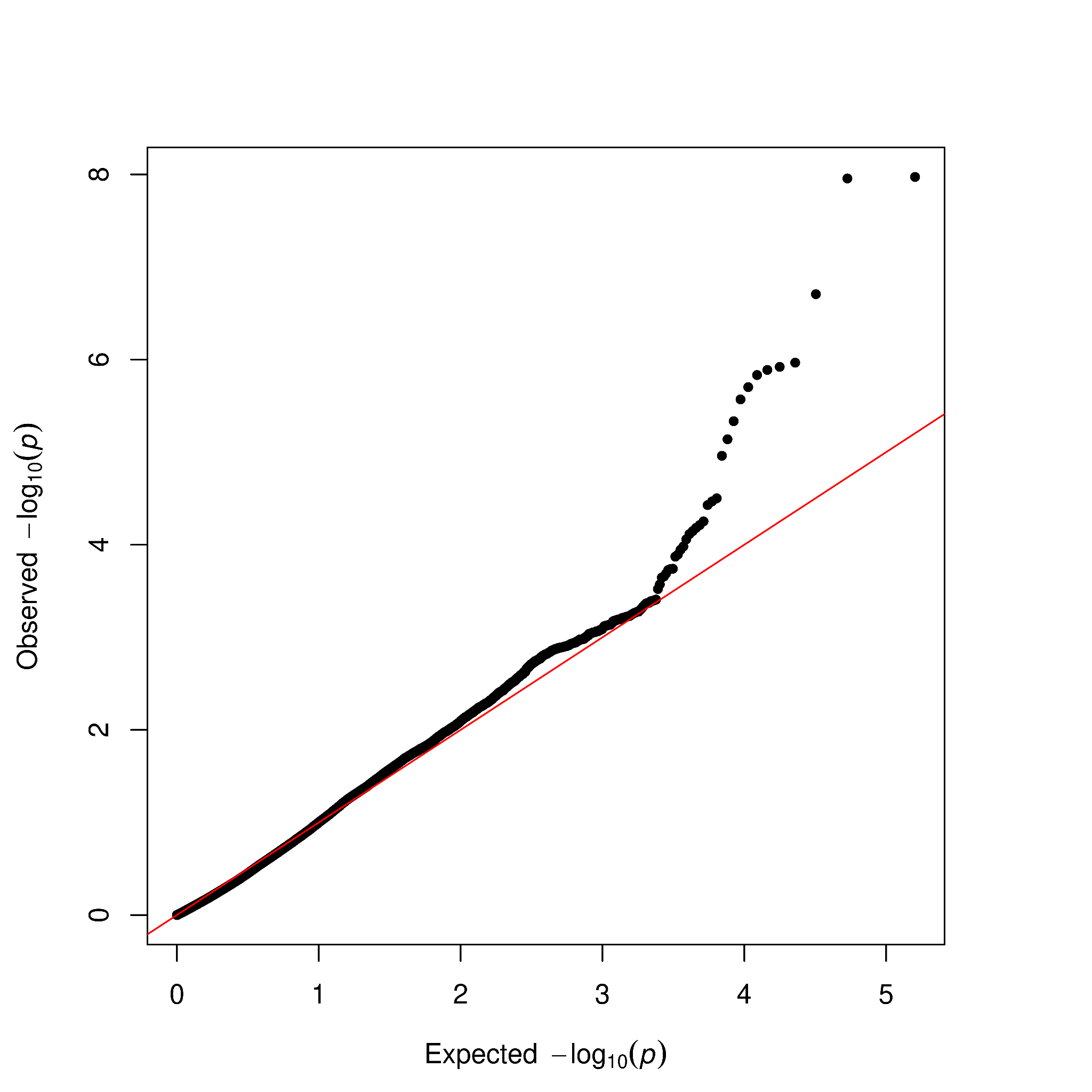

Figure S19. Q-Q plot for GLMM-based GWAS for body height (large individuals).

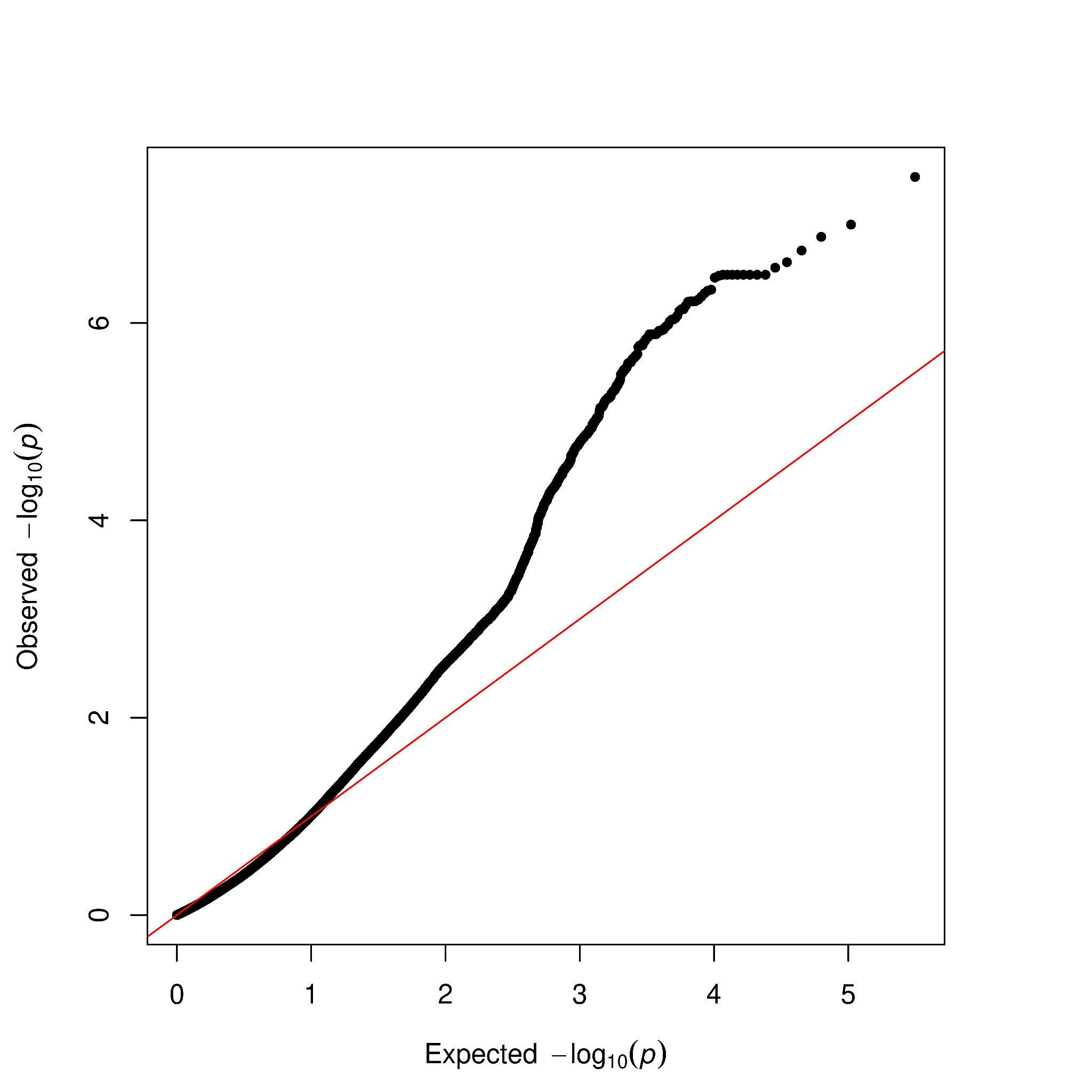

Figure S20. Q-Q plot for GLMM-based GWAS for body weight (all individuals).

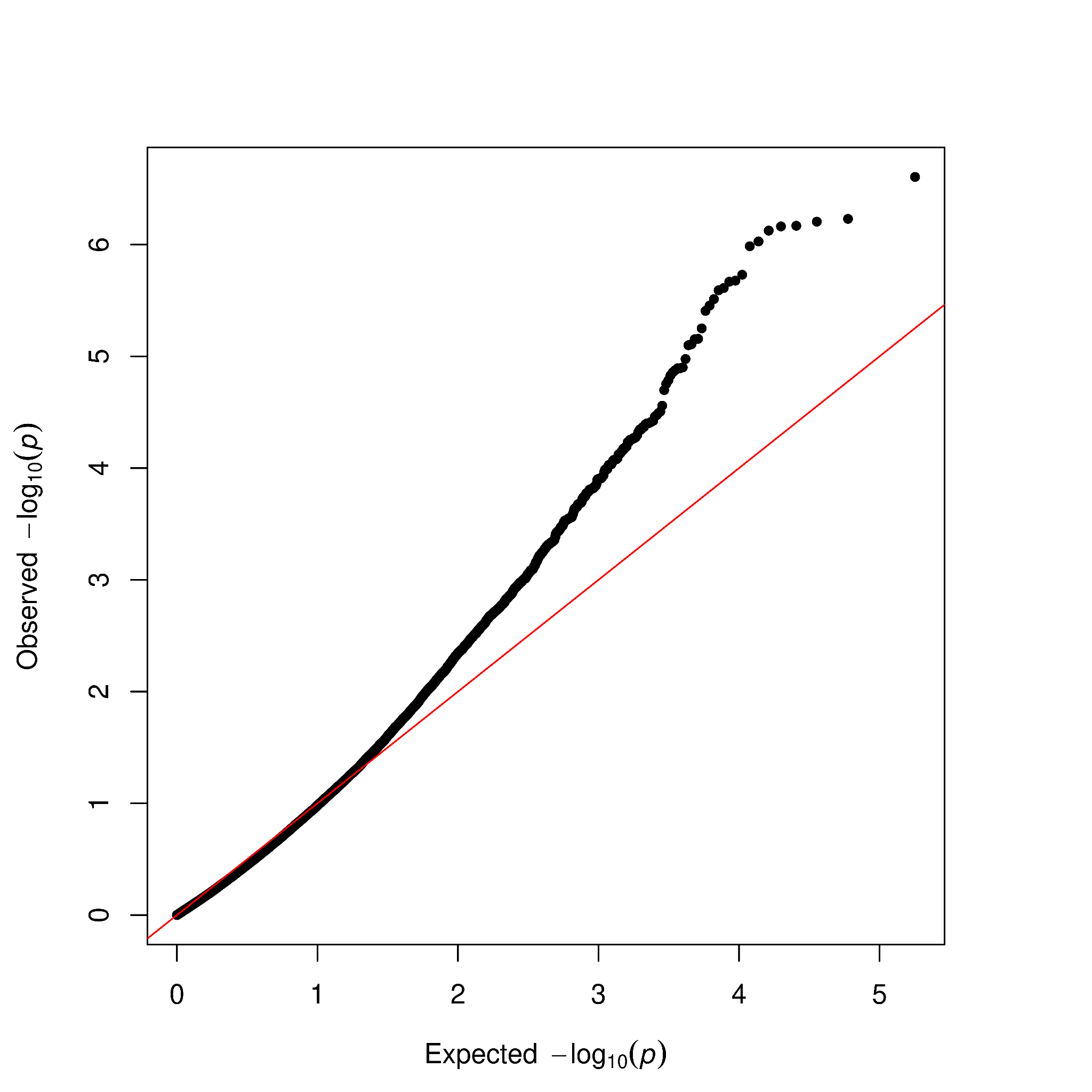

Figure S21. Q-Q plot for GLMM-based GWAS for body weight (small individuals).

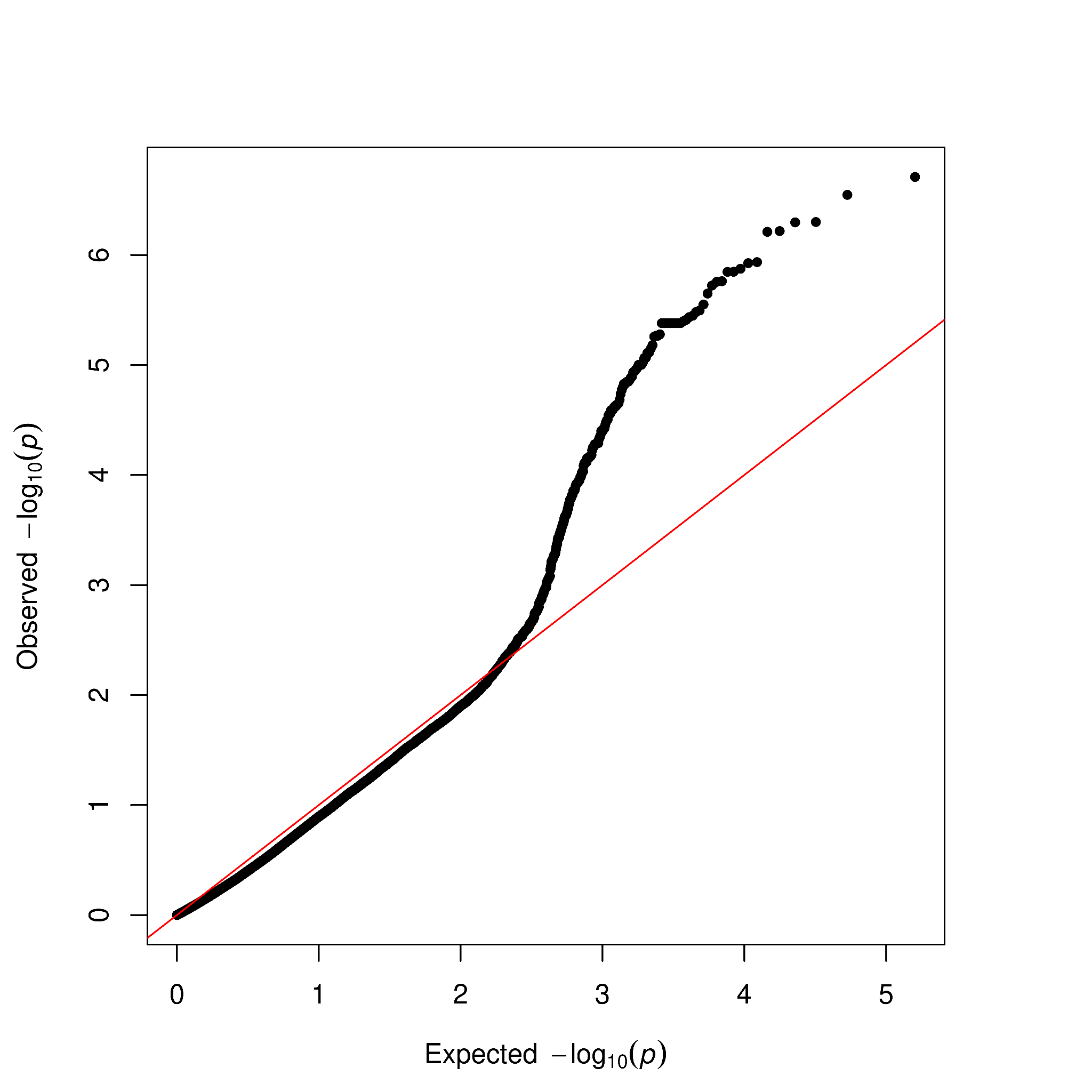

Figure S22. Q-Q plot for GLMM-based GWAS for body weight (large individuals).

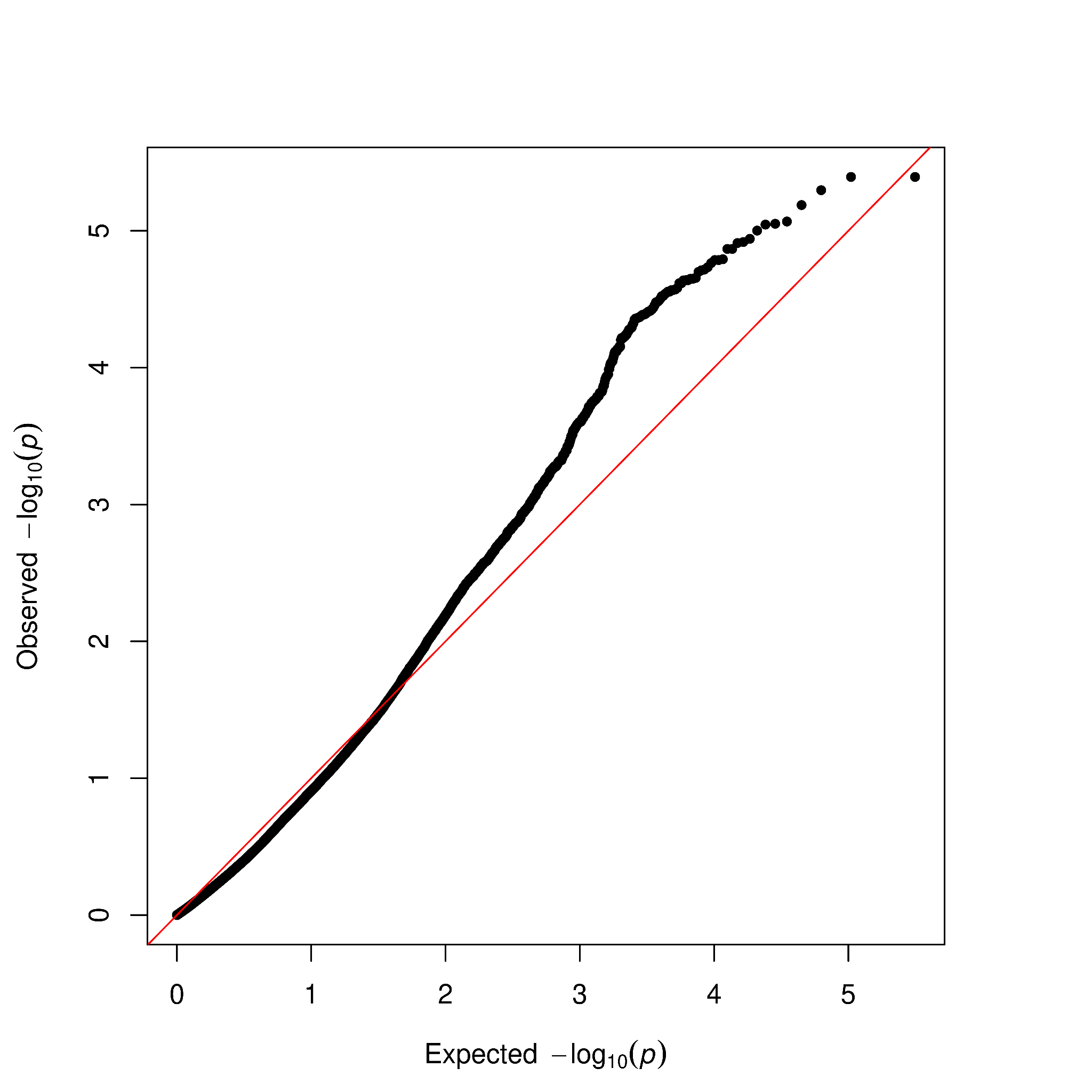

Figure S23. Q-Q plot for GLMM-based GWAS for lifespan (all individuals).

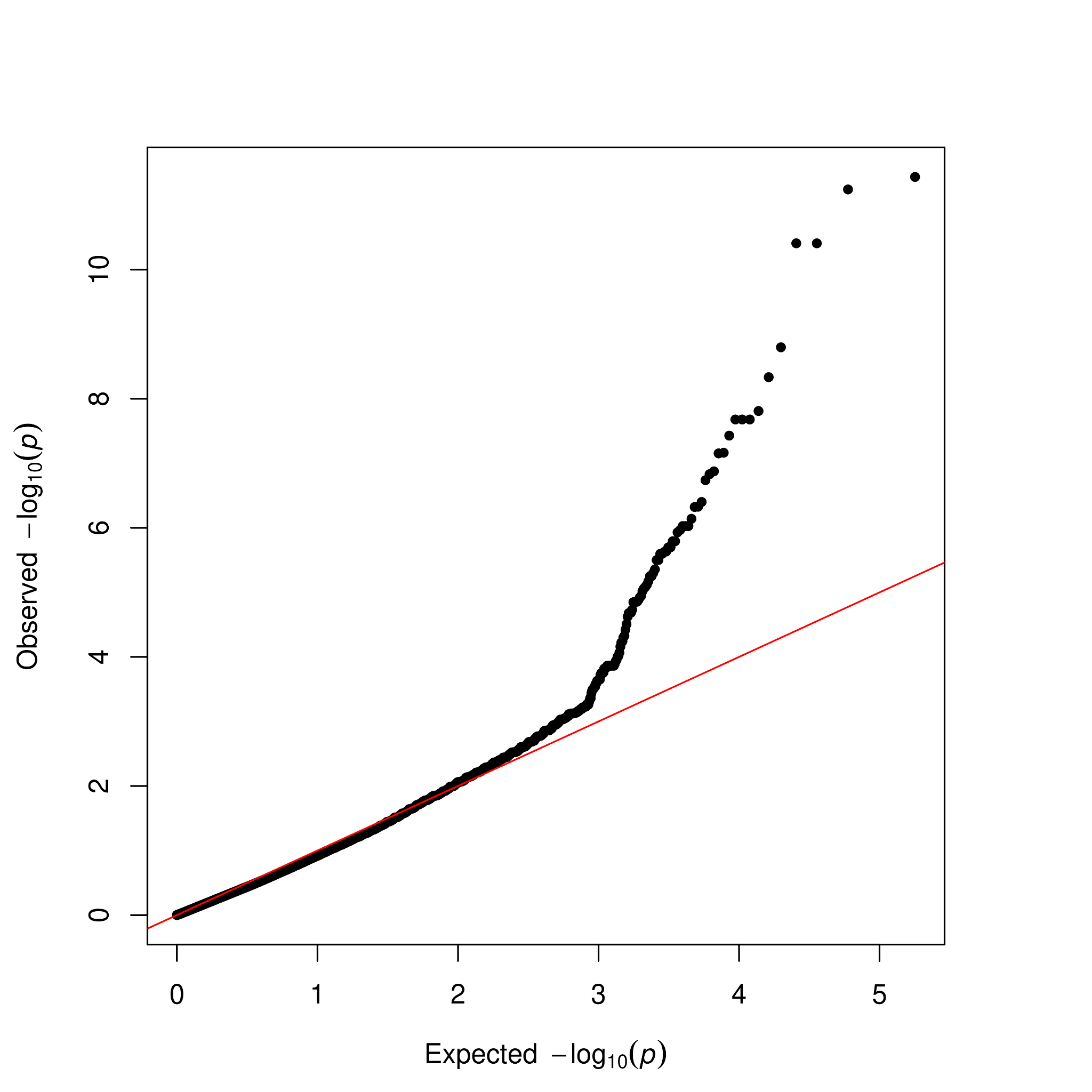

Figure S24. Q-Q plot for GLMM-based GWAS for lifespan (small individuals).

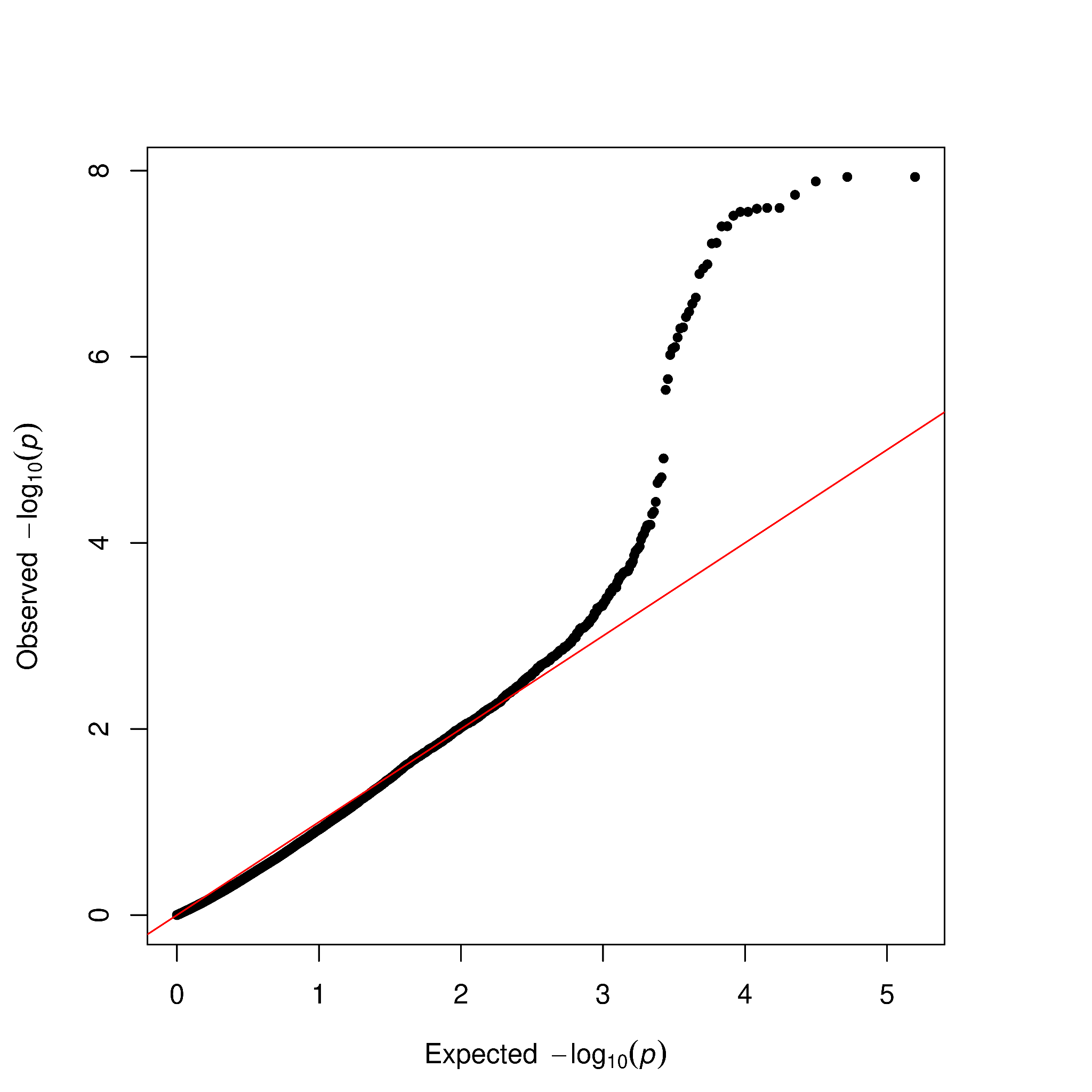

Figure S25. Q-Q plot for GLMM-based GWAS for lifespan (large individuals).

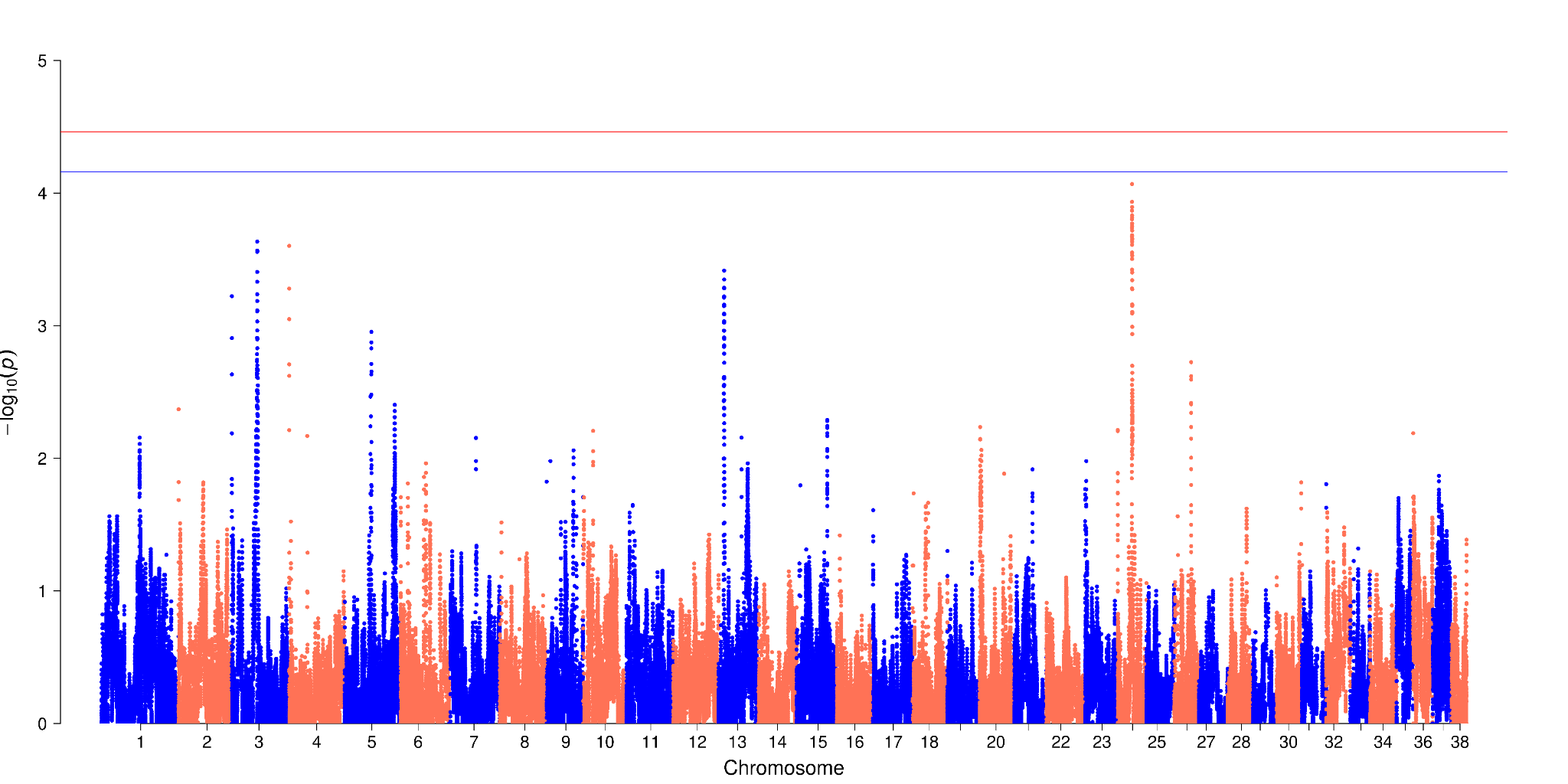

Figure S26. Manhattan plots for GLMM-based GWAS using presence/absence of ROH for furnish. The x-axis represents the genomic position. The y-axis represents the log10 base transformed p-values. Single nucleotide polymorphisms are represented by a single point. The red horizontal line indicates the genome wide significance (GWS) threshold and the blue horizontal line indicates the suggestive wide significance (SWS) threshold. The average marker density for each chromosome can be found in Supplementary Table 16.

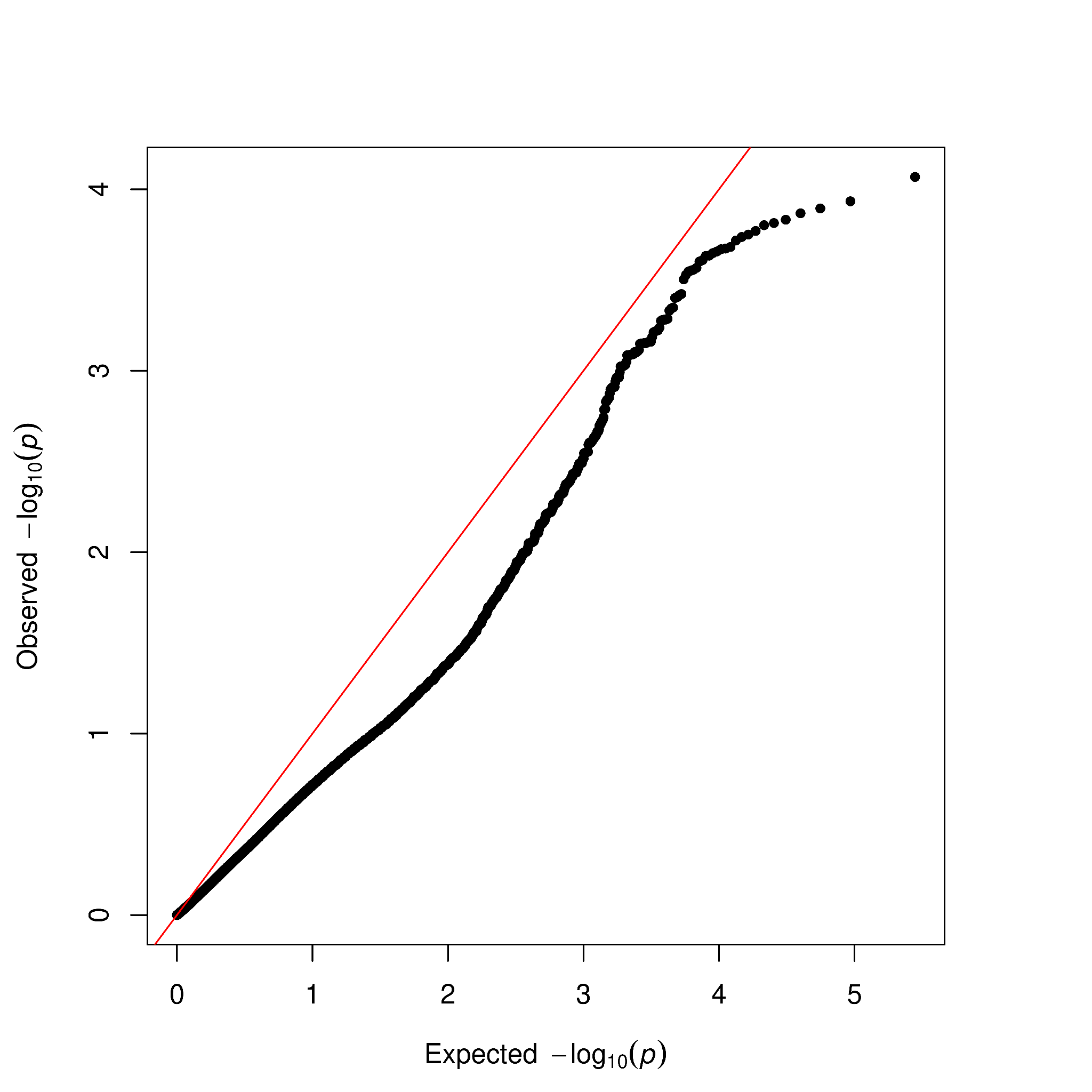

Figure S27. Q-Q plot for GLMM-based GWAS for Furnish.

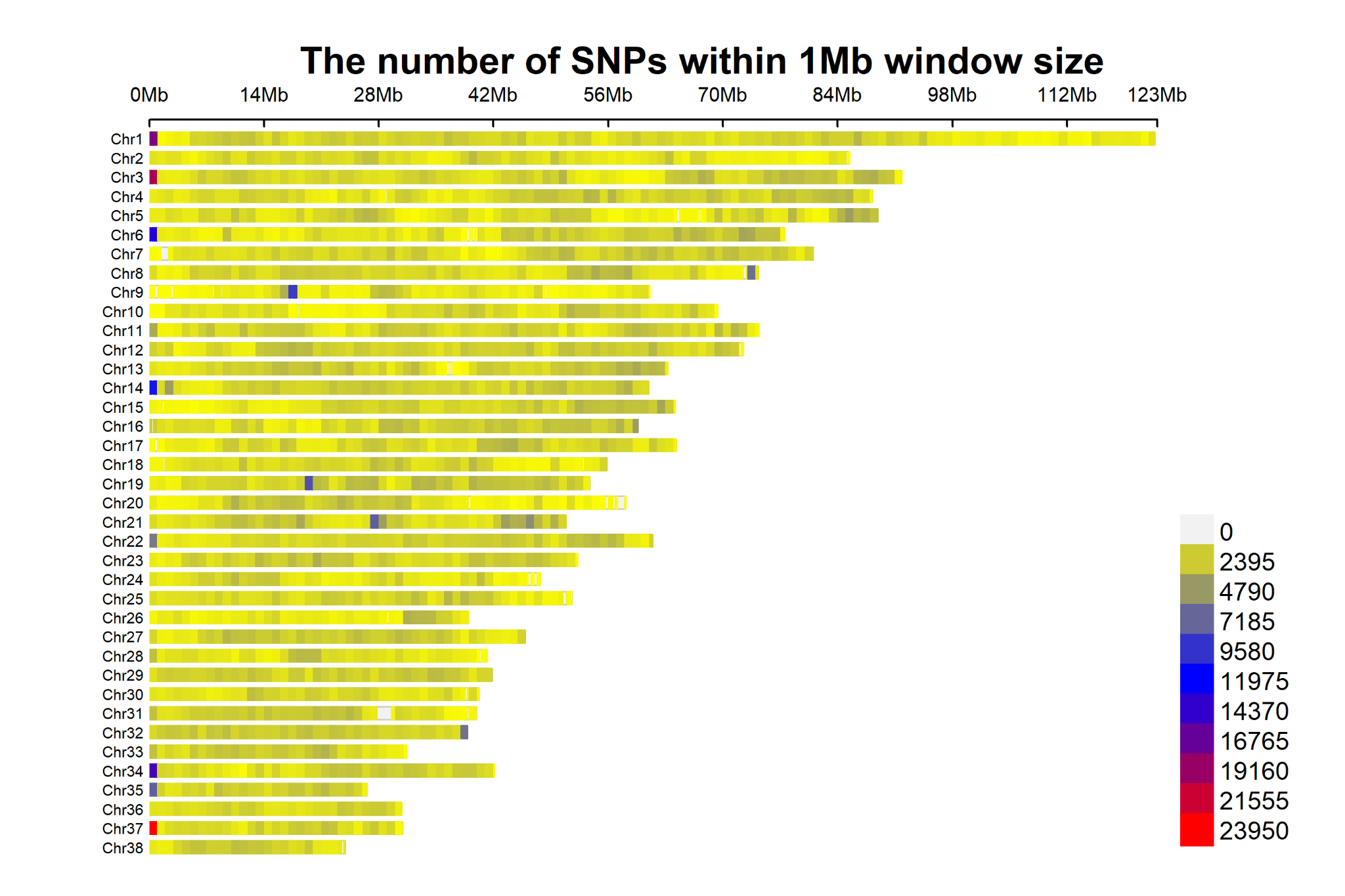

Figure S28. SNP density plot across 38 chromosomes representing the number of SNPs within 1 MB window size. The x-axis represents the chromosome length in Mb. The y-axis represents the chromosome number. Different colors correspond to SNP density.

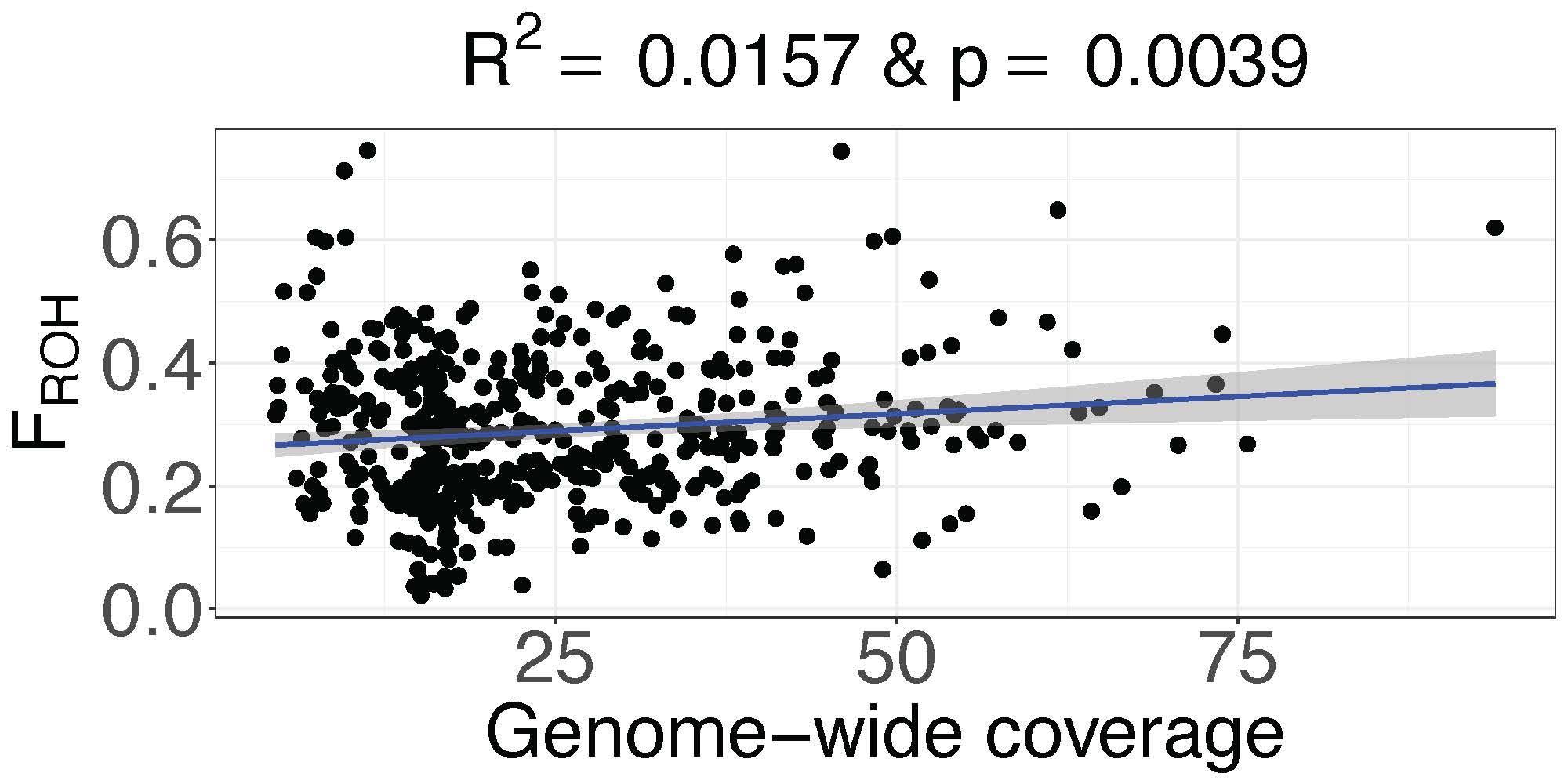

Figure S29. Relationship between genome wide coverage and FROH for all 466 individuals. The x-axis represents genome wide coverage. The y-axis represents the FROH. Different colors correspond to SNP density. The blue line represents the fitted regression line, and the gray line is the 95% confidence interval for the linear model.

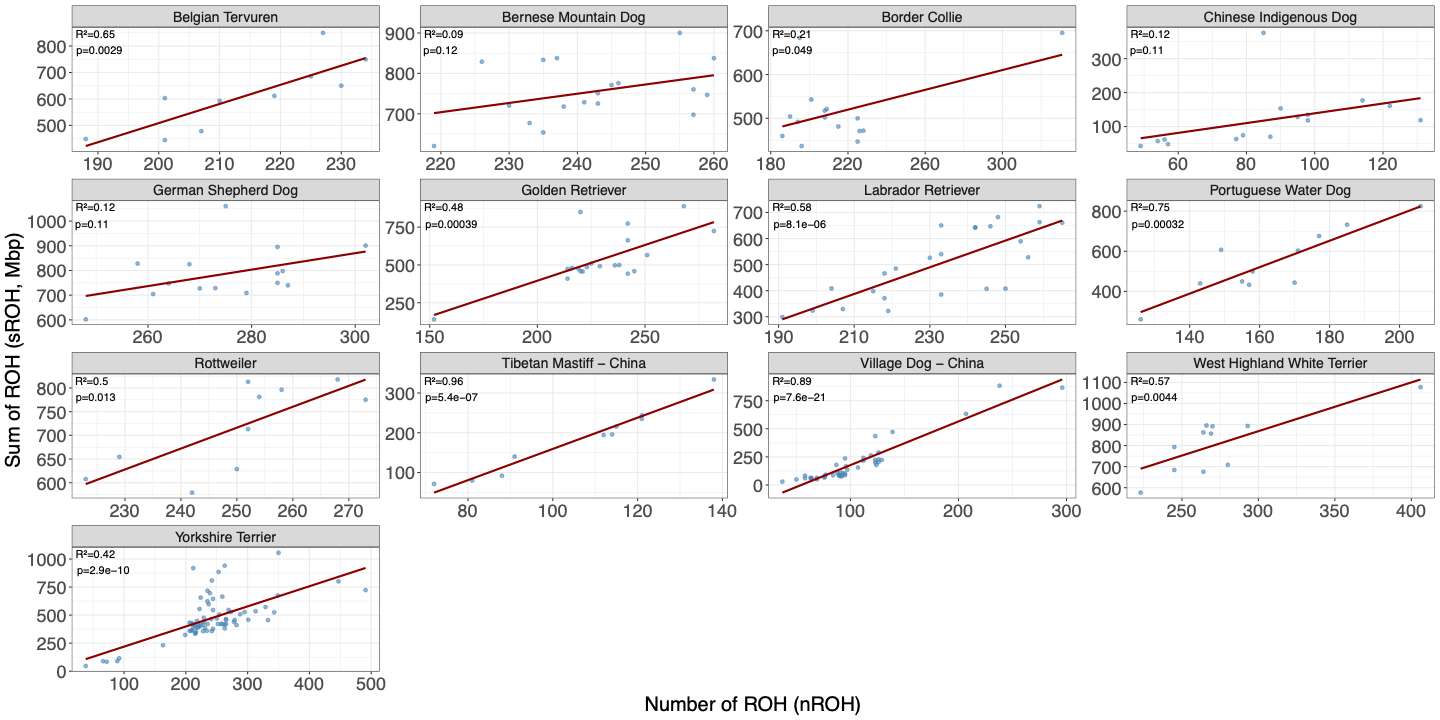

Figure S30. Relationship between nROH and sROH for all groups with more than 10 individuals. Adjusted R^2^ and p-value are reported for each group.

Figure S31. Clade average effect-size computed across 1000 simulations for lifespan using individual level data (blue) and breed average beta (orange) and empirical values of F_ROH_.

### Supplementary Text

Assume we have $N$ breeds $b_{1}, \ldots, b_{N}$ and each breed has $M_{i}$ dogs, so we have $M_{1}, \ldots, M_{N}$.

Suppose that the trait score of each breed is normally distributed with $N(\mu_{i}, \sigma_{i}^{2})$, i.i.d. and

F_ROH_ is normally distributed with $N(0, \sigma_{i}^{2})$ across all breeds.

Proof:

Given data ($X_{ij_{i}}, Y_{ij_{i}}$), where:

- $i$ : breed ∈ $1,\ldots,N$
- $j_{i}$ : dog index ∈ $1, \ldots, M_{i}$ where $M_{i}$ is $\#$ of dogs in breed $i$.
- $Y_{ij_{i}} \sim N\left( \mu_{i}, \sigma_{i}^{2} \right),$ where $\sigma_{i}^{2}$ is small for all $i$, is the trait score of each breed.
- $X_{ij_{i}} \sim N(0, \sigma^{2})$ is the F_ROH_ for all individuals.

We want to show that $Y = \beta_{1}X + \beta_{0}$ (regression from ($X_{ij_{i}}, Y_{ij_{i}}$)) is close to $Y = \alpha_{1}X + \alpha_{0}$ (regression from ($X_{ij_{i}}, Y_{ij_{i}}$)).

To demonstrate that $\beta_{1}$ is sufficiently close to $\alpha_{1}$ we can use MLE regression.

By MLE regression:

- $\beta=\frac{Cov(X_{ij_{i}}, Y_{ij_{i}})}{Var(X_{ij_{i}})}$
- $\alpha=\frac{Cov(X_{ij_{i}}, \mu_{i})}{Var(X_{ij_{i}})}$

Thus, we only need to show $Cov(X_{ij_{i}}, Y_{ij_{i}})$ is close to $Cov(X_{ij_{i}}, \mu_{i})$ for all i.

Because $Y_{ij_{i}}= \mu_{i}+N(0, \sigma_{i}^{2})$

$Cov\left( X_{ij_{i}}, Y_{ij_{i}} \right)=Cov(X_{ij_{i}}, \mu_{i}+N(0, \sigma_{i}^{2}))$

By Law of Total Covariance:

$$Cov\left( X_{ij_{i}}, Y_{ij_{i}} \right)=Cov(E\left( X_{ij_{i}} | i \right), E(Y_{ij_{i}} | i))+ E(Cov\left( X_{ij_{i}}, Y_{ij_{i}} \right| i))$$

$$=Cov\left( X_{ij_{i}}, \mu_{i} \right)+ Cov(X_{ij_{i}}, \mu_{i}+N(0, \sigma_{i}^{2}))$$

$$=Cov\left( X_{ij_{i}}, \mu_{i} \right)+ Cov(X_{ij_{i}}, N\left( 0, \sigma_{i}^{2} \right))$$

$$\approx Cov\left( X_{ij_{i}}, \mu_{i} \right)$$
